# Large inversions in Lake Malawi cichlids are associated with habitat preference, lineage, and sex determination

**DOI:** 10.1101/2024.10.28.620687

**Authors:** Nikesh M. Kumar, Taylor L. Cooper, Thomas D. Kocher, J. Todd Streelman, Patrick T. McGrath

## Abstract

Chromosomal inversions are an important class of genetic variation that link multiple alleles together into a single inherited block that can have important e7ects on fitness. To study the role of large inversions in the massive evolutionary radiation of Lake Malawi cichlids, we used long-read technologies to identify four single and two tandem inversions that span half of each respective chromosome, and which together encompass over 10% of the genome. Each inversion is fixed in one of the two states within the seven major ecogroups, suggesting they played a role in the separation of the major lake lineages into specific lake habitats. One exception is within the benthic sub-radiation, where both inverted and non-inverted alleles continue to segregate within the group. The evolutionary histories of three of the six inversions suggest they transferred from the pelagic Diplotaxodon group into benthic ancestors at the time the benthic sub-radiation was seeded. The remaining three inversions are found in a subset of benthic species living in deep waters. We show that some of these inversions are used as XY sex-determination systems but are also likely limited to a subset of total lake species. Our work suggests that inversions have been under both sexual and natural selection in Lake Malawi cichlids and that they will be important to understanding how this adaptive radiation evolved.

## INTRODUCTION

Large genomic inversions are a particularly interesting type of genetic variation in the evolutionary process^1–4^. These rearrangements of DNA sequence strongly suppress recombination between inverted and non-inverted alleles, enabling the capture and accumulation of genetic variants of smaller molecular size that are linked together on each inversion haplotype^5^. These alternative haplotypes largely follow independent evolutionary paths within a species, accumulating new mutations that can affect phenotypes under natural and sexual selection. Structural rearrangements can be large, spanning multiple megabases of DNA containing hundreds of genes^2,6^. They have been shown to play a role in adaptive divergence within a variety of species^7–12^, in creating alternative reproductive strategies and mating types^13–16^, in evolution of sex chromosomes^1,17^, and in the formation of pre- and post-zygotic barriers between incipient species^18,19^.

When comparing the genomes of extant species, chromosomal inversions are often found as fixed differences, consistent with a role in speciation^20–22^. Inversions have been shown to play a role in adaptation to different environments and the establishment of prezygotic and postzygotic barriers^21,23^. However, the evolutionary forces responsible for establishing and spreading inversions remain obscure. Natural selection and sexual selection are thought to play a role but neutral forces like genetic drift can also be responsible for their spread. Perhaps most likely, all of these different evolutionary forces contribute to their fixation^24,25^.

Few vertebrates offer as much potential for investigating evolution and speciation as the *Cichlidae* family of fish, one of the most species-rich and diverse families of vertebrates, with an estimated 2,000-3,000 species across the globe ^26–28^. Evolutionary radiations have occurred at least three times in the African Great Lakes (Lake Malawi, Lake Tanganyika, and Lake Victoria). In Lake Malawi, more than 800 species of cichlids evolved in the past 1.2 million years^27,29–31^. Speciation in cichlid fishes involved extremely high levels of phenotypic divergence, including changes in body shape, jaw structures, and feeding behaviors related to prey in their ecological niches^26,32^. Additionally, traits that create prezygotic barriers between species are also extremely diverse. Color patterns are quite dramatic and diverse among species and include both differences in color and species-specific patterns including bars, stripes, and blotches that are thought to reinforce barriers between species in sympatry^33^. In some species, mating is seasonal, with gatherings on ‘leks’ where males build bowers by scooping and spitting out sand over the course of a week or more^34^. Females choose mates based upon features of these bowers including shape and size^35^. Most of the Lake Malawi species can be induced to interbreed in the lab, providing the opportunity to use genetic approaches to study causality of associated changes. Despite the presence of these prezygotic barriers, substantial gene flow has occurred multiple times within the lake^30,36^.

Our current understanding of how the Lake Malawi radiation created >800 species is a set of three serial diversifications, starting from a single riverine-like ancestor lineage, that created seven major ecogroups that currently live in the lake^30^ (**Figure 1**). Each ecogroup forms its own clade in a whole-lake phylogeny and occupies distinct habitats within Lake Malawi. A pelagic lineage separated first, further diversifying into a deep-water ecogroup (*Diplotaxodon*) and a mid-water ecogroup (*Rhamphochromis*). A muddy/sandy benthic lineage evolved from the ancestral riverine lineage next, which split into three ecogroups (collectively referred to as *benthics*) living either in the water column (*utaka*), over shallow sandy/muddy shores (*shallow benthics*), or in deep-water habitats (*deep benthics*). Finally, an ecogroup living over rocky habitats evolved (*mbuna*). The riverine generalist *Astatotilapia calliptera (AC)* living in the border regions of the lake and surrounding rivers, is the seventh ecogroup, and is thought to represent the ancestral lineage that seeded the three primary Lake Malawi lineages. Subsequent radiations within these ecogroups based on trophic specializations and sexual selection further diversified the species flock^30,32^.

**Figure 1.**
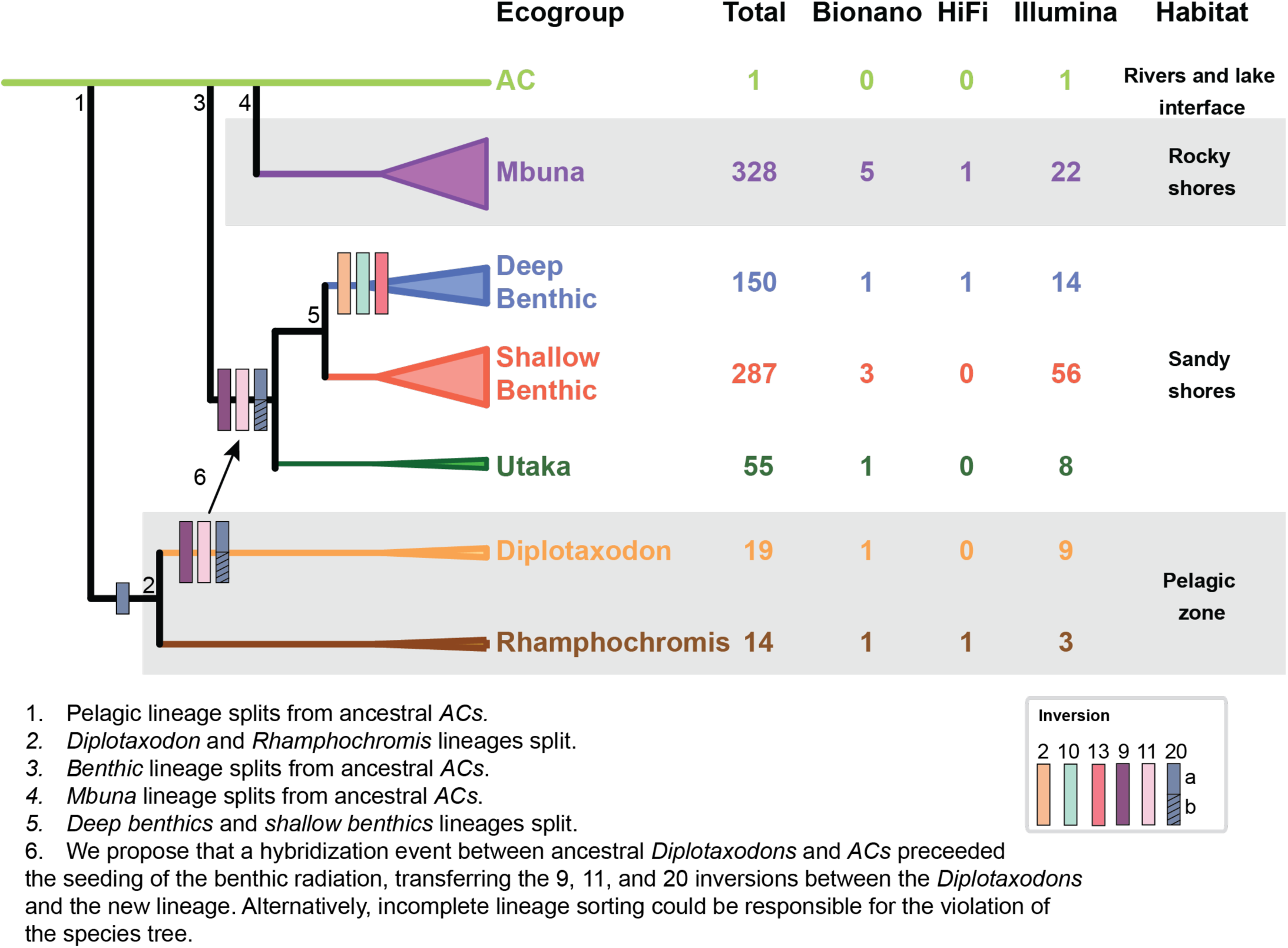
Overview of Lake Malawi cichlids. A summary of the Lake Malawi cichlids and the results from this paper. On the left, a phylogeny showing the seven major lineages that currently live in the lake which are referred to as ecogroups. AC is *Astatotilapia calliptera*, a riverine species that lives in the margins of the river/lake interface. For each ecogroup, we summarize the 1) estimated number of species in the lake, 2) the number of species we have Bionano optical read data for, 3) the number of species we have PacBio based genome assembies, 4) the number of species with Illumina short-read sequencing data, and 5) the primary habitat these ecogroups live in. Deep and shallow benthics are distinguished by the lake bottom depths where they live while utakas live in the water column but mate over the shore. We also summarize the distribution of 6 large inversions we have identified, colored by the chromosome they are found on (legend). We propose that the inversions on 9, 11, and 20 originated in the pelagic lineage and spread to the benthics by hybridization very early on in the evolution of the benthic lineage. Alternatively, these inversions distribution could be explained by incomplete lineage sorting.

Here, we investigate the role large inversions played in the Lake Malawi radiation using three technologies. Using the Bionano Saphyr optical mapping system, we identify four single large inversions and two double inversions, ranging from 9.9Mb to 20.6Mb in size. A fifth single inversion, composed of just one of the two tandem inversions on chromosome 20, was also identified, indicating a serial set of rearrangements created this structural variant. We validate these inversions using new genome assemblies created using PacBio Hifi reads. Finally, we use existing and newly generated short-read Illumina sequencing to genotype these inversions in DNA samples from cichlids collected from Lake Malawi. These inversions primarily segregate by ecogroup, with no inversions found in the *mbuna* and *AC* samples, one inversion fixed in *Rhamphochromis*, three inversions fixed in *Diplotaxodon*, and all six segregating within the *benthic* sub-radiation. The evolutionary histories of these inversions are inconsistent with the species phylogeny and suggest that the three *Diplotaxodon* inversions spread into the *benthics* via hybridization around the time the benthic lineage diverged from the ancestral riverine species. The three additional inversions are found primarily in *benthic* species living in deep water habitats. We provide evidence that three of the inversions are involved in sex determination in a subset of *benthic* species. Our work provides a framework for understanding the role of inversions in the adaptive radiation of Lake Malawi cichlids, and supports future investigations into their roles in the establishment of ecogroups, adaptation to deep water habitats, and in controlling traits under sexual selection.

## RESULTS

### Improvement of the Metriaclima zebra reference

The *mbuna* cichlid, *Metriaclima zebra*, was the first species sequenced as a reference genome for Lake Malawi cichlids^37^. The current version of its genome, M_zebra_UMD2a, has been created by assembling PacBio reads into contigs which were anchored to 22 chromosomes using recombinant maps generated using crosses between different Lake Malawi species^38^. While the overall quality of this reference is high compared to other non-model organisms, a total of 20.5% of the DNA was not anchored to a chromosome. Additionally, the orientation of contigs can be ambiguous given how contigs were anchored to linkage groups using recombinant maps. These imperfections could complicate inversion discovery, so we first sought to improve the reference sequence.

The Bionano Saphyr system generates large (150 kb - multiple Mbs) DNA molecules and labels a specific motif (CTTAAG) with a fluorescent probe^39^. While these molecules do not provide sequencing information, the distance between observed fluorescent loci can be used to assemble multiple molecules into large maps, with lengths spanning up to an entire chromosome. Comparison of these maps with an *in-silico* digest of the reference sequence identifies large structural changes in the observed sample or errors in the reference genome. We processed DNA collected from the blood of three *Metriaclima zebra* individuals using the Bionano pipeline and assembled DNA molecules into optical maps. We identified many discrepancies between the observed optical maps and the M_zebra_UMD2a assembly (**Figure 2**). These differences fall into two primary categories: 1) contigs that were incorrectly ordered or oriented and 2) unplaced contigs that Bionano anchored to specific chromosomes. The first category is consistent with errors in the assembly caused by ambiguity in placing contigs into a linkage group using recombination maps. Our results indicate that 178 of the contigs are incorrectly oriented and 133 of the unanchored contigs could be placed onto specific chromosomes.

**Figure 2.**
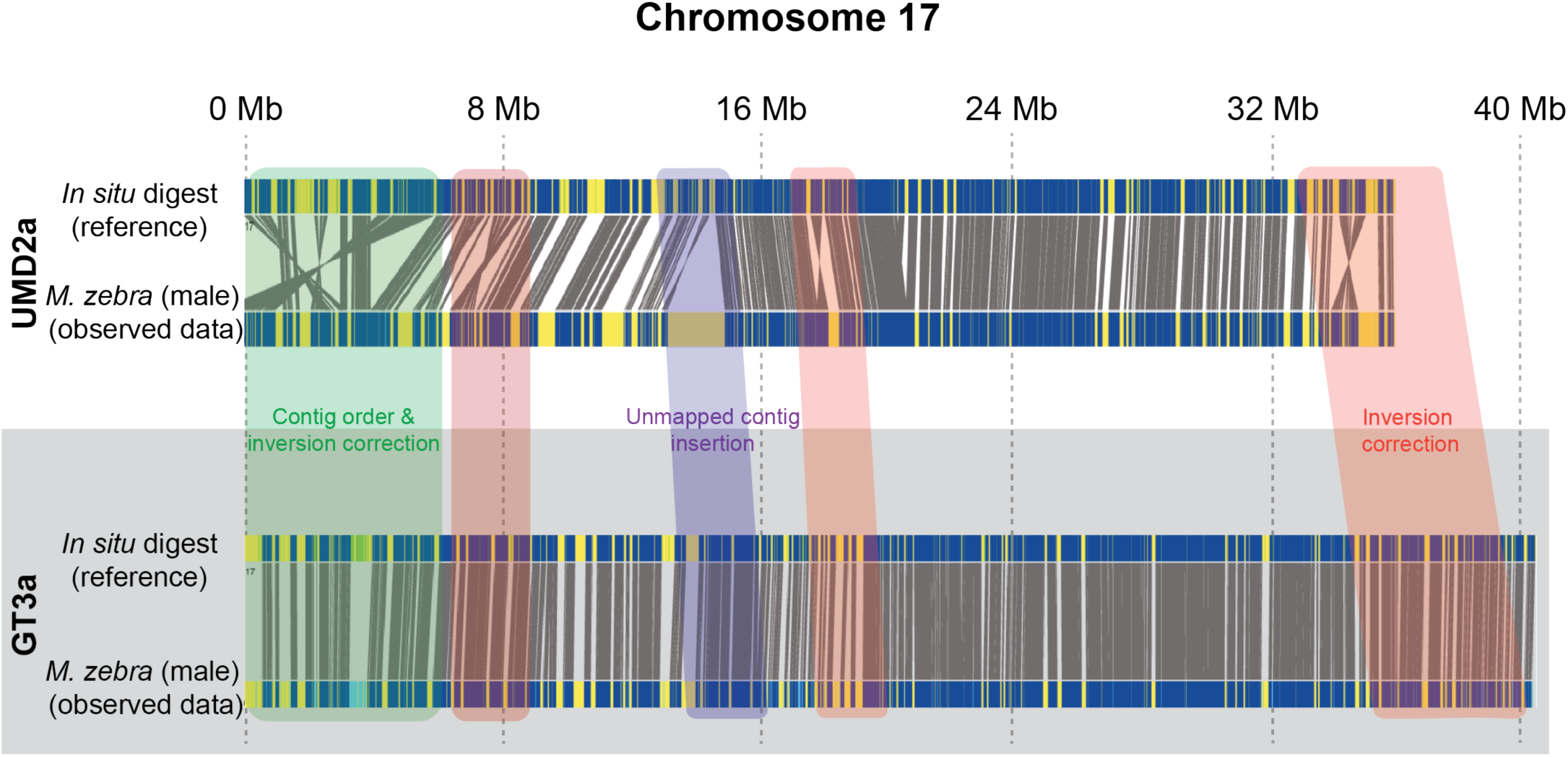
Generation of a new high quality reference genome for *Metriaclima zebra*. Alignment of Bionano maps derived from a single *Metriaclima zebra* male to two different reference sequences for Chromosome 17 (white and grey, respectively). M_zebra_UMD2a reference (white) is the existing reference genome and M_zebra_GT3a reference (grey) was created for this manuscript using a combination of PacBio HiFi reads and Bionano molecules. For each reference, the top bar shows the expected location of each CTTAAG motif (blue) created from an *in silico* annotation of the reference sequence. These are aligned (grey lines) to observed locations in maps generated from Bionano Saphyr optical molecules (bottom bar). Deviations from co-linearity indicate reference errors (green, blue, and red highlights). Yellow indicates regions of the genome that lack aligned motifs. A summary of the improvements in the new genome is found in **Table S1**.

To further improve the reference, we used PacBio HiFi reads generated from an additional male *Metriaclima zebra* individual to create a new genome assembly, taking advantage of the low overall error rate (< 0.1%) of HiFi reads compared to previous generation PacBio technologies. Contigs generated from these data were combined with Bionano maps to generate a new hybrid genome assembly, which we refer to as the M_zebra_GT3a assembly. The overall quality of the assembly was excellent, with 933Mb of 962 Mb of DNA assigned to the 22 linkage groups, (**Table S1**) reducing the amount of unplaced contigs from 196 Mb to 28 Mb. The N50 length was 32.542Mb, with many of the scaffolds containing entire chromosomes.

### Identification of six large inversions from 11 species of Lake Malawi cichlids

Armed with the improved reference, we used the Bionano system to test 31 samples from 11 different species found in Lake Malawi. Eight of the species were bred in the Streelman or McGrath laboratories (**Table 1 and Table S2**). These included four *mbuna* species (*Pseudotropheus demasoni*, *Cynotilapia zebroides*, *Labeotropheus fuelleborni, and Labeotropheus trewavasae),* two *shallow benthic* species (*Mchenga conophoros* and *Nyassachromis prostoma* “Orange Cap”), a single *deep benthic* species (*Aulonocara sp. ‘chitande type north’ Nkhata Bay*), and a single *utaka* species (*Copadichromis virginalis)*. To expand our coverage of the Lake Malawi radiation, we also obtained samples from the pet trade for a single species of the *Diplotaxodon* ecogroup, *Diplotaxodon limnothrissa,* and a single species of the *Rhamphochromis* ecogroup, *Rhamphochromis longiceps*. Finally, we also tested a single male *Protomelas taeniolatus shallow benthic* individual collected from Lake Malawi based on its position in principal component analysis of SNVs described below. In total, we were able to obtain samples from 6 of the 7 ecogroups, excluding *Astatotilapia calliptera*.

**Table 1.**
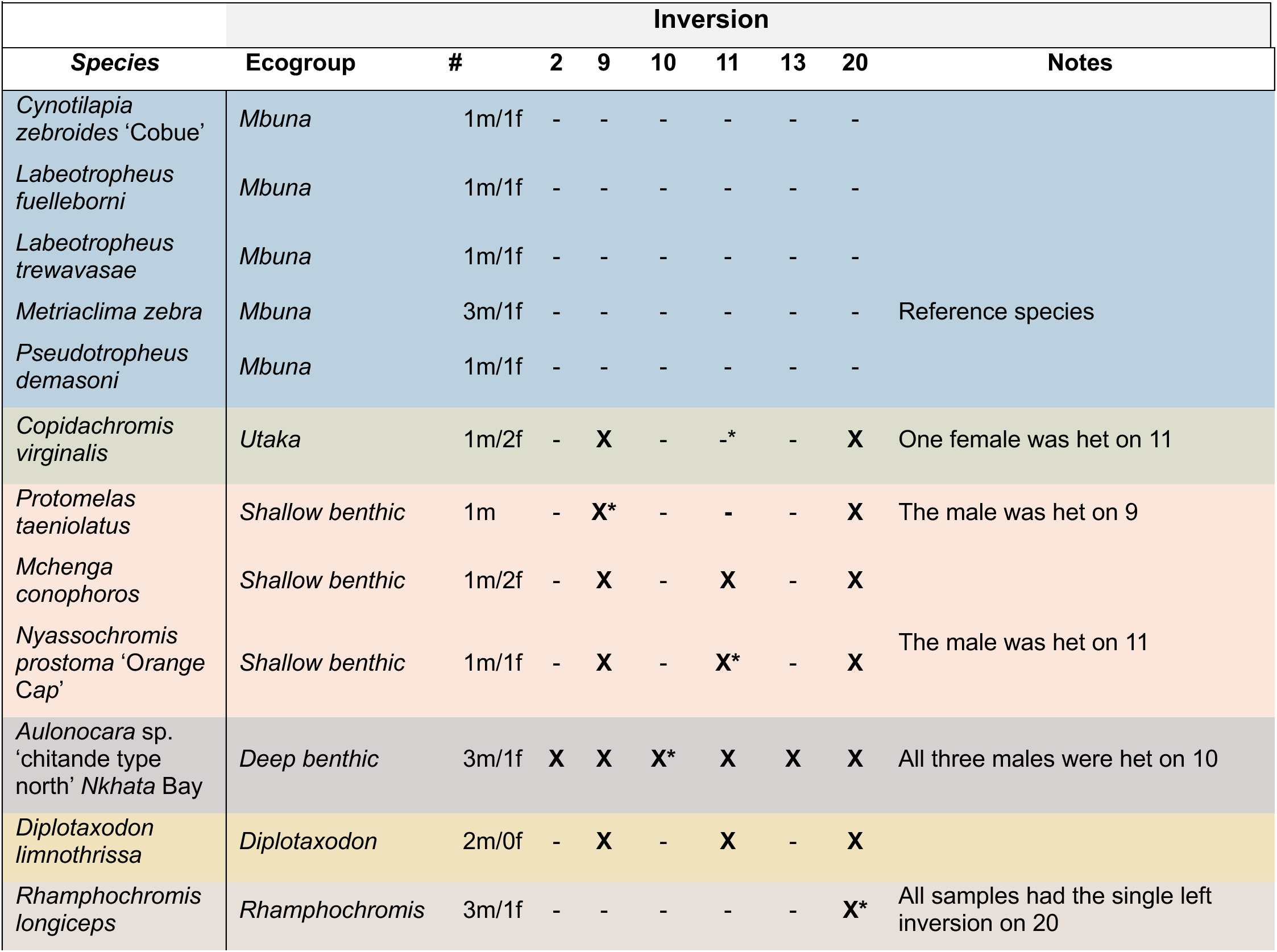
Inversion genotypes of 31 samples based on Bionano data.

From these 31 individuals, we identified six new inversions ranging from 9.9 - 22.9 Mb in size (**Figure 3**): single inversions on chromosomes 2, 9, 10, and 13, and double inversions on chromosomes 11 and 20. Interestingly, regarding the double inversion on chromosome 20, we also identified four *Rhamphochromis* individuals that carried only the first inversion, lacking the 4.1 Mb long second part of this rearrangement (**Figure 3**). This indicates that the tandem inversions on 20 were formed by at least a two-part process, most parsimoniously with the left arm of the inversion (20a) forming first followed by the right inversion (20b) occurring in that genetic background.

**Figure 3.**
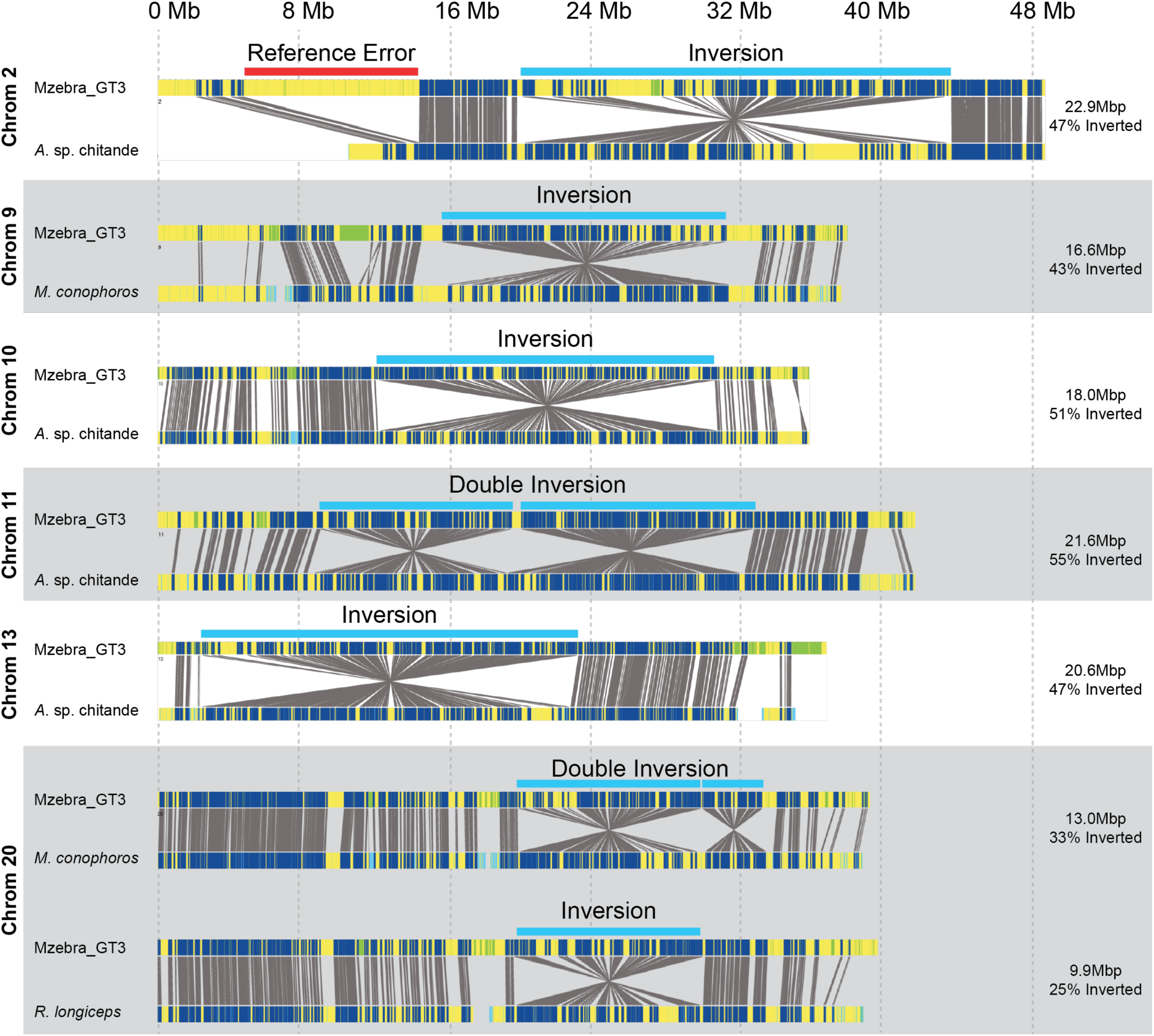
Identification of six large single or double inversions segregating within Lake Malawi cichlids. Representative alignments of inversions or double inversions identified from blood samples obtained from 30 individuals from eleven species. For each inversion, the top shows predicted CTTAAG motifs from an *in-silico annotation* of the reference genome, the bottom shows motifs identified using Bionano molecules obtained from an individual of the species indicated on the left, and the grey lines indicate matching motifs based upon predicted and observed distances. Yellow indicates regions containing no matches between motifs in the reference genome and the mapped sample. Single inversions were identified on chromosomes 2, 9, 10, 13, and 20. Tandem double inversions were identified on chromosomes 11 and 20. The estimated length and percentage of the chromosome spanned by the inversion is shown on the right. The single inversion on 20 identified in *Rhamphochromis longiceps* has the same position as the first inversion of the double inversion on 20 identified from *Mchenga conophoros*. A list of all samples and their inversion genotype is in Table 1. Note that an error or polymorphism in the reference genome in chromosome 2 is indicated in the first panel.

The distribution within these species is summarized in **Table 1**. For simplicity, we will refer to all six rearrangements as inversions, regardless of whether they are single or double inversions. The most common inversions were on chromosome 9, 11, and 20 which were found in a combination of *benthic* and *Diplotaxodon* individuals. The position of these three inversions were also consistent with regions of low recombination rate previously identified in an intercross between a *mbuna* and *benthic* individual^38^, as expected, since inversions suppress recombination between inverted and non-inverted haplotypes. The remaining three inversions (2, 10, and 13) were only found in the deep benthic individuals (*Aulonocara sp. ‘chitande type north’ Nkhata Bay*). While for most samples the inversion genotype was homozygous, for five individuals the optical maps supported both inverted and non-inverted haplotypes, indicating the inversions were heterozygous in those samples (**Figure S1**).

### *de novo* reference genomes support the presence of these six inversions

To provide further support for these structural rearrangements, we generated *de novo* hybrid genome assemblies of three *Aulonocara sp. ‘chitande type north’ Nkhata Bay* individuals, which carry all six of the identified inversions, and two additional *Metriaclima zebra* individuals using a combination of PacBio HiFi reads and Bionano maps. The overall quality of these assemblies is high, with large N50 values and a small number of individual contigs (**Table S3**). We performed whole genome alignments between the five new

assemblies and the GT3a reference genome (**Figure S2-S7**). These alignments show that the new assemblies support the presence of all six inversions in the *Aulonocara sp. ‘chitande type north’ Nkhata Bay* individuals. For most inversions, a single scaffold covered the entire inversion with breakpoints in locations consistent with the Bionano maps. The two new control assemblies were consistent with the M_zebra_GT3a reference and did not show any evidence of inversions in those regions. In other regions of the genome, however, there were differences in the alignment of the M_zebra_GT3a assembly and the *Metriaclima zebra* genomes, most prominently in the left arm of chromosome 2 (**Figure 3**) and in the highly repetitive chromosome 3, which likely indicates some errors in the M_zebra_GT3a assembly or that a large polymorphism segregates within the species.

We refined the location of the end points for each inversion using the genomic alignments. However, the presence of a large amount of repetitive DNA in regions near these regions prevented us from defining inversion boundaries at single base pair resolution. The mechanisms that create large inversions often involve repetitive sequence near the breakpoints, making exact localization difficult if not impossible^40^.

We also took advantage of a genome assembly for a *Rhamphochromis sp ‘chilingali’* individual (fRhaChi2.1) created by the Darwin Tree of Life Project produced using PacBio data and Arima2 Hi-C data (Accession PRJEB72870). The genome alignment between the fRhaChi2.1 assembly and the GT3a assembly confirmed a single inversion on chromosome 20, consistent with the Bionano data (**Figure S2- S7)**.

For each inversion, we determined the ancestral haplotype using two species as outgroups, *Oreochromis niloticus* and *Pundamilia nyererei. Oreochromis niloticus,* or Nile tilapia, is an important food fish that’s estimated to have diverged from Lake Malawi cichlids 14.1 to 30 million years ago while *Pundamilia nyererei* is a more closely related haplochromine found in Lake Victoria^37,41^. For this analysis we used a published whole genome assembly for *Oreochromis niloticus* (UMD_NMBU)^38^ and generated a hybrid genome assembly for *Pundamilia nyererei* using a previously published whole genome assembly^37^ combined with Bionano maps we produced from a single individual (**Figure S8**). For all six inversions, the reference haplotype found in the GT3a assembly represents the ancestral state.

### Inference of the segregation of these inversions in Lake Malawi using short-read sequencing

The distribution of these inversions within the lake can provide important information to their function. Genotyping inversions using short read sequencing is possible because each haplotype is on an independent evolutionary trajectory due to the suppression of recombination. Population genetics approaches take advantage of the fact that inversions create patterns of inheritance between SNVs that are readily detected by approaches such as Principal Component Analysis (PCA)^42^.

We analyzed short read sequencing data from the 31 samples with Bionano data along with 297 wild individuals published by the Durbin lab^30^, 15 wild individuals published by the Streelman lab ^31,43^, and 22 wild individuals sequenced for this paper, aligning reads and calling small variants against our new M_zebra_GT3a reference genome. PCA analysis was used to analyze the major axes of variation across the whole genome and within the six inversions. We separately analyzed the left and right inversions on 11 and 20. For the whole genome, the species within an ecogroup clustered together within the first two principal components, reflecting their evolutionary history. This matched the previously reported results of Malinsky et al^30^. Individuals from four of the ecogroups - *Rhamphochromis*, *Diplotaxodon*, *Astatotilapia callipter*a, and *mbuna* - formed four distinct clusters (**Figure 4A and Figure S9**). A fifth, more diffuse cluster, composed of *shallow benthics*, *deep benthics*, and *utakas*, was also present. The broader distribution of the *benthic* ecogroups in PC space is consistent with both their recent separation as well as large amounts of gene flow thought to have occurred between these different species^30,36^.

**Figure 4.**
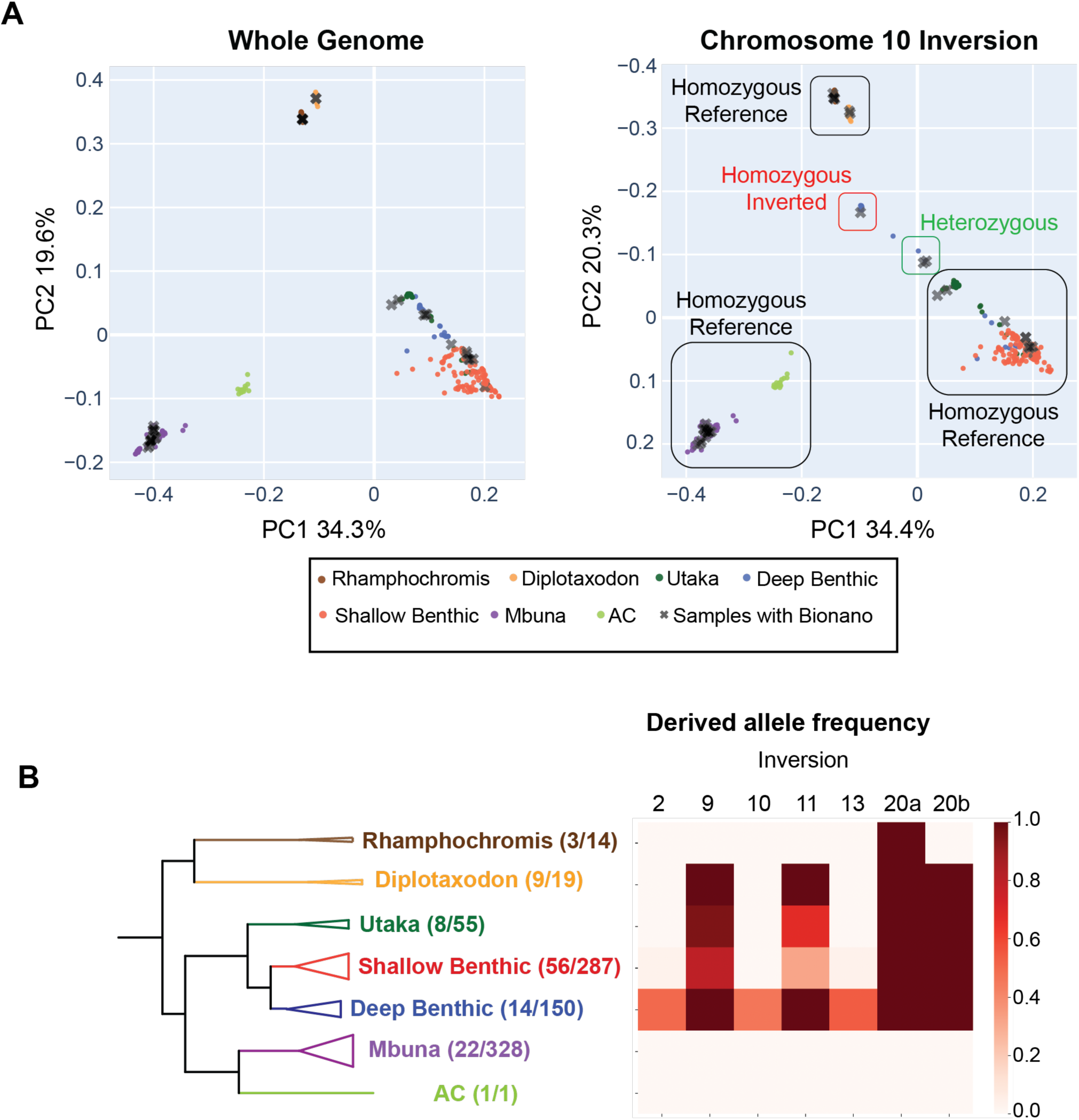
Distribution of inversions within the Lake Malawi ecogroups. Principal component analysis was used to analyze SNVs distributions in 365 samples within each inversion to genotype the inversion haplotype for all six inversions. (**A**) We illustrate the approach for genotyping the chromosome 10 inversions. The PCA plot for the entire genome is shown on the left, with each sample colored by its ecogroup. Samples from each ecogroup cluster together, reflecting their evolutionary history. Individuals with Bionano data are shown as grey Xs. On the right, for the SNVs that fall within the chromosome 10 inversion, the deep benthic/shallow benthic and utaka individuals split into multiple clusters. Using the samples genotyped with Bionano data, we assign each cluster to an inversion genotype, which is shown by the black, green, and red boxes. To make comparisons of the whole genome PCA and chromosome 10 PCA plots easier, we reversed the y-axis on the right panel. The assigned genotypes of each sample can be found in Supplemental Table 2. Interactive PCA plots for each inversion are included in **Figures S9-S17**. (**B**) The derived allele frequency was calculated for each inversion within the seven ecogroups. The number of species that were genotyped and the estimated number of species that live in Lake Malawi are listed in the whole-genome phylogeny (left).

The PCA plots using the SNVs within the six inversions often violated the pattern of the whole genome PCA (**Figure 4A** and **Figure S10-S17**). Using samples that were sequenced with both Illumina and Bionano technologies, we assigned each cluster to specific inversion genotypes (**Table S2**). We did not have Bionano data for two clusters (one on chromosome 9 and one on 11 – see **Figures S11, S13, and S14**), so we supplemented our dataset by adding Bionano data from an additional *shallow benthic* individual (*Protomelas taeniolatus*) that fell into both clusters. The PCA clusters varied in their complexities for each inversion depending largely on their distribution within the *benthic* ecogroup (more on this below) and included both homozygous genotypes and heterozygous genotypes. We summarize the overall distribution of the inversions within each of the ecogroups in **Figure 4B**.

All specimens of the *mbuna* and *AC* ecogroup were fixed for the homozygous non-inverted genotype for all six inversions. In *Rhamphochromis*, all the inversions were homozygous non-inverted except for the left arm of 20 which was fixed for the single inversion state. The *Diplotaxodon* individuals all carried the non-inverted alleles for the inversions on 2, 10 and 13 and were fixed for the inversions on 9, 11, and 20.

The distribution of these inversions was the most complicated within the *benthics*. The inversions on 2, 10, and 13, which were identified in *Aulonocara sp. ‘chitande type north’ Nkhata Bay*, were mostly found together in other *deep benthics*, including the *Alticorpus* (*geo7reyi*, *macrocleithrum*, *peterdaviesi*), *Aulonocara* (*blue chilumba*, *gold* and *minutus)*, and *Lethrinops* (*gossei, longimanus ‘redhead’, and sp. olivera*) genera. The inversions on 2 and 13 were also found in the shallow benthic species *Placidochromis longimanus*. However, not all the *deep benthics* carried the three inversions, including five *Aulonocara* species (*baenschi*, *gertrudae*, *steveni*, *stuartgranti*, and *stuartgranti* Maisoni) and a single *Lethrinops* species (*longipinnis*). The distribution of these three inversions in *benthic* species known to inhabit the deepest habitats are suggestive for a role in depth adaptation.

Both inversions on 20 were homozygous inverted in all benthic individuals. For the single inversion on 9, many *benthics* were homozygous for the inverted haplotype. However, an additional PC cluster was present, composed of 64 *shallow benthic* and *utaka* individuals heterozygous for the non-inverted haplotype (**Figure S1**). Interestingly, no *benthic* individuals were identified as homozygous for the non-inverted state.

Finally, the distribution of the chromosome 11 inversion was also determined. Individuals of all three genotypes (homozygous non-inverted, homozygous inverted, and heterozygous) were observed (**Figure 4a**).

### Evolutionary history of 9,11, and 20 suggest introgression from Diplotaxodon into benthic ancestors

The distribution of these inversions conflicts with the species tree, suggesting that these inversions could have spread to the *benthics* via hybridization or carried forward by incomplete lineage sorting. To determine their evolutionary history, we built phylogenies for the whole genome as well as for each inversion using SNVs that fell within each of the six inversions using representative species from each ecogroup (73 total – **Table S2**) as well as two previously sequenced *Pundamilia nyererei* individuals as an outgroup (**Figure 5A** and **Figures S18 – S25**)^44,45^. To avoid issues with haplotype phasing, we focused on individuals that were homozygous for one of the two inverted genotypes.

**Figure 5.**
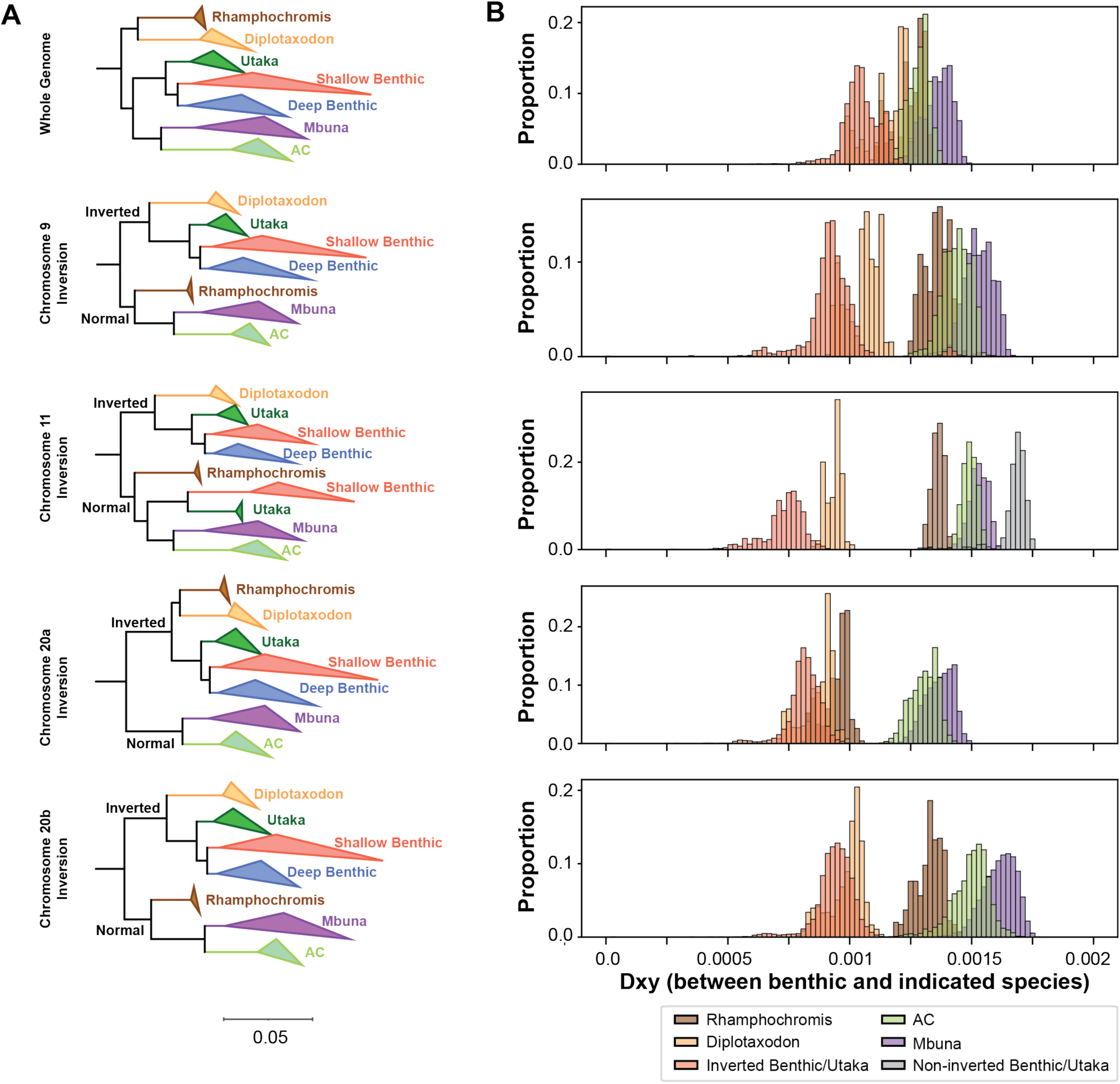
Evolutionary history of large inversions violates species tree. (**A**) Maximum likelihood trees built from either whole genome SNVs (top) or within the 9, 11, and 20 inversions. Between 2-25 representative samples were selected for each ecogroup (see **Table S2**) and individuals that were heterozygous were excluded from this analysis. Clades were collapsed by ecogroup for visualization purposes. Full phylogenies are presented in **Figures S18** to **S25**. The whole genome tree indicates *Diplotaxodon* and *Rhamphochromis* species split first and formed their own clade. For each of the displayed inversions, the first split was between samples carrying the inverted vs. non-inverted genotypes. Additionally, the *Diplotaxodon* individuals now formed a clade with the *benthics* that carry the inversion. (**B**) Density plots of *d_XY_* values comparing *benthic* species and species of the indicated ecogroup. For the displayed inversions, the *Diplotaxodon* species were much closer in evolutionary distance when compared to the whole genome plot.

While the whole genome phylogeny matched the expected topology based upon our understanding of the Lake Malawi species tree, the topologies of each of the inversions showed key differences between the separation of ecogroups. The *benthic* group often split into separate clades, correlated with their inversion genotype. Prominently, we found the three inversions on 9, 11, and 20 showed key differences in the relationship between the *Diplotaxodon* and *benthic* ecogroups. While *Diplotaxodon* is a sister group to *Rhamphochromis* at the species level, for these three inversions, the *Diplotaxodon* individuals formed a clade with the *benthics* that also carried the inversion. Additionally, genetic distance between *Diplotaxodon* and *benthics* carrying the inversion was also much smaller than the rest of the genome (**Figure 5B**). Both the phylogenies and genetic distances between ecogroups are consistent with the introgression of the 9, 11, and 20 inversions from the *Diplotaxodon* to *benthic* ancestors at the time of the benthic radiation.

### A role for large inversions in sex determination within the benthic ecogroup

The evolution of sex determination is often associated with the presence of inversions to repress recombination between the sex chromosomes^46,47^. We tested whether these inversions could play a role in sex determination in *benthic* species, as we identified individuals that were heterozygous for some of these inversions (**Table 1**). We started with the inversion on 10, as our Bionano data identified three male *Aulonocara sp. ‘chitande type north’ Nkhata Bay* individuals that were heterozygous and a single female that was homozygous for the inverted haplotype. To test whether this inversion is linked to a genetic locus (or loci) that act as an XY system in this species, we tested two separate broods (Brood 1; 2 parents, 24 offspring and Brood 2: 2 parents, 24 offspring) currently growing in the laboratory, sequencing 12 males and 12 females from each brood with Illumina paired-end sequencing. We used PCA to genotype each of the 52 animals (4 parents and 48 offspring) for the chromosome 10 inversion genotype. The association in the offspring between the inversion 10 genotype and sex was almost perfect: 24 females and 1 male were homozygous for the inversion, while 23 males were heterozygous for the inversion (**Figure 6B**). These individuals were sexed by venting so the imperfect segregation could either indicate an error in sexing or that other modifiers influence sex.

**Figure 6.**
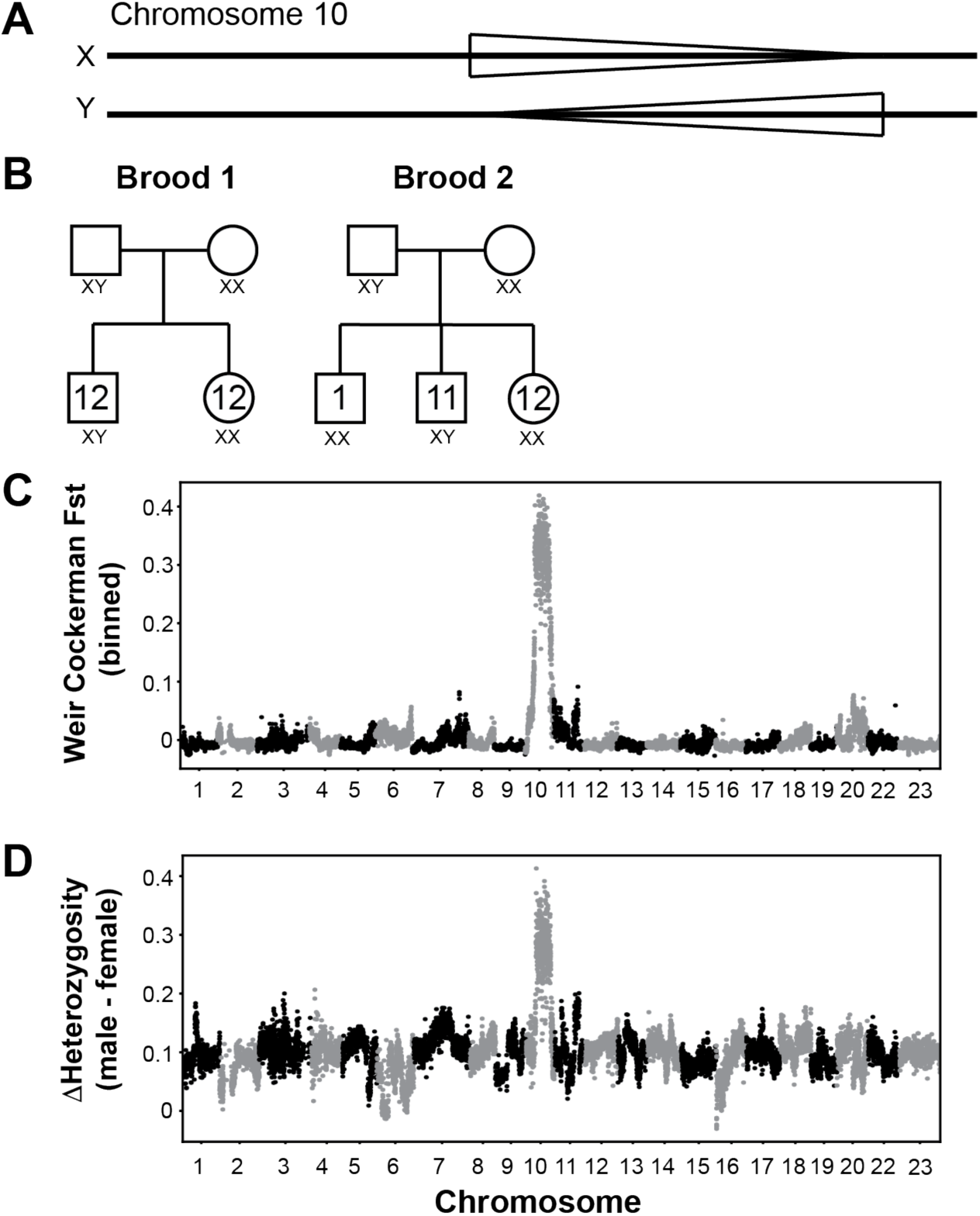
A role for the chromosome 10 inversion in sex determination. (**A**) Schematic of the two inversion states (normal and inverted) and their assignment as X or Y chromosomes. (**B**) 24 offspring (12 male and 12 female) from two separate male/female pairs were genotyped for the inversion. For 47 of 48 individuals, the XY animals were male and the XX animals were female. One male was XX. (**C**) F_ST_ analysis comparing 24 males and 24 females identified chromosome 10 as the primary genetic region that segregated with sex. (**D**) Comparison of heterozygosity levels of 24 male and 24 female animals also identified chromosome 10 as an outlier.

We also used binned Weir Cockerham F_st_ analysis, grouping male and female individuals to scan the genome for regions that were associated with sex (**Figure 6C**). The inversion on 10 was the region of the genome that was in strongest linkage disequilibrium with sex. Males also had a much higher heterozygosity in this region than females (**Figure 6D**). Altogether, these results indicate that the inversion on 10 acts as an XY sex determiner in the *Aulonocara sp. ‘chitande type north’ Nkhata Bay* species.

We also tested eight laboratory *Nyassachromis prostoma* “Orange Cap” individuals, as our Bionano data identified a male that was heterozygous for the inversion on 11 and a female that was homozygous for the inversion. We performed short-read sequencing on three males and five females growing in the lab, genotyping the inversion using PCA analysis. All three males were heterozygous for the inversion while the five females were homozygous, a significant association between genotype and sex (Fisher exact test p- value = 0.0179). This suggests a separate XY system segregates in *Nyassachromis prostoma* “Orange Cap”, linked to the inversion on chromosome 11.

## DISCUSSION

Here, we characterize the presence of large inversions found in the Lake Malawi cichlid flock, identifying six inversions segregating within the lake, providing a framework to address the role structural variants played in this lake’s adaptive radiation (**Figure 6**). What are the evolutionary forces responsible for their spread and what role do they currently play in the lake? Inversions have multiple, well-characterized roles in adaptive evolution: 1) they can capture beneficial alleles responsible for adaptation to local habitats, 2) they can capture alleles that create prezygotic and postzygotic barriers that prevent gene flow during speciation, and 3) they can capture alleles that influence sex-specific phenotypes that are linked to sex-determination systems^1,3,4^. These inversions could have also spread through neutral processes like genetic drift or through other adaptive processes not listed here.

This question is much more complicated in the context of the Lake Malawi cichlid radiation with ∼800 different species. The role of these inversions must be considered in a species- and time-specific way. The recent history of the lake was quite dynamic, with large cyclical changes in water depth and shoreline in the past 800,000 years that are predicted to create alternating periods of extinctions and radiations^48–50^. Similarly, the role the inversions play in each species (such as sex determination) will not be the same. Determining whether these inversions carry alleles that contribute to the phenotypic diversity found in Lake Malawi will be an important first step towards resolving the evolutionary forces responsible for their spread through the lake. This is possible as most species in the lake can interbreed, allowing for quantitative genetics approaches to link genotype variation to phenotypic variation and determine what traits are influenced by the inversions.

The local adaptation mechanism is one well-characterized method to explain how inversions can spread through adaptive evolution^6,8^. Under this hypothesis, two populations undergoing adaptive divergence in the presence of migration and gene flow create conditions that can select for inversions, provided they capture and link adaptive alleles together that increase fitness to their local environments^51^. Apart from the *benthics*, the inversions are fixed in each ecogroup in the inverted or non-inverted state, each of which live in unique habitats in the lake that place very different evolutionary pressures on the animals that live within them. In this scenario, the inversions would capture adaptive alleles that increase fitness of ancestral species to pelagic and deep-water environments. To illustrate this scenario, we consider the time the pelagic groups split from the ancestral AC fish that is thought to live in rivers and the borders of the lake (**Figure 1**). If the first inversion on 20 captured alleles that allowed these pelagic ancestors to inhabit the lake, it could be selected on provided that these pelagic ancestors continue to interbreed with the AC ancestors. Similarly, the diversification of the *Rhamphochromis* and *Diplotaxodon* groups could have involved the inversions on 9, 11, and right arm of 20, and the adaptation of *benthics* to deeper water habitats could have involved the inversions on 2, 10, and 13. We do note that the existing theory for recruiting inversions to local adaptation does not account for multiple unlinked inversions (i.e. 9, 11, and right arm of 20). Future experimental and comparative genomics approaches can determine if local adaptation was at play at the time these lineages separated.

We also provide evidence that two inversions play a role in sex determination in the *benthics*, with the strongest evidence showing the inversion on 10 acts as an XY system in a deep benthic species. Additionally we show that the inversion on 11 is significantly associated with sex determination in a shallow benthic species as an XY system. Despite the importance of the maintenance of distinct, reproductively compatible sexes, sex determination systems can be evolutionarily labile, and Lake Malawi cichlids show exceptional variability, with at least 5 different genetic sex determination systems described to date^52,53^. While we implicated two inversions in sex determination, our work also underscores how dynamic sex determination is within the *benthics*. While many benthic *species* carry the different inversions, their genotypic distribution in most species is inconsistent with a role for sex determination. Effectively, the genetic determination of sex determination largely must be determined in a species by species way, and individual species may be segregating multiple sex determination systems. The selective forces on sex chromosomes are strong, including balancing selection for sex ratio and sexually antagonistic selection on sexually dimorphic phenotypes. Lake Malawi cichlids show amazing sex differences, characterized by diversification in numerous phenotypes, including pigmentation, behavior, and morphology thought to contribute to species barriers. The use of inversions in sex-determination allows for the accumulation of alleles in linkage with the sex-determiner gene which will have sex-specific effects in the male individuals that carry the Y chromosomes. The presence of these inversions in various ecogroups could simply be remnants of their role in sex determination in the ancestors of pelagic ecogroups, and have nothing to do with local adaptation, which could have contributed to their spread before sex chromosome turnover ended balancing selection on the inverted/non-inverted haplotypes and subsequent fixation of these inversions. Identification of the causal alleles for sex determination on these inversions will help us to resolve this question.

The presence of non-inverted haplotypes in the *benthics* could have been retained from the time the lineage initially formed, with both haplotypes under balancing selection if they were used for sex determination. A recent preprint, however, which also identified these 6 inversions presented here using multiple technologies, also utilized short-read sequencing of 1375 individuals to create an alternative model^54^. They inferred the non-inverted haplotypes of the 9 and 11 inversions were introgressed back into the *benthics* from riverine species within and outside of the lake. Similarly, the non-inverted haplotypes on chromosomes 2, 10, and 13 in the *deep benthics* were introgressed from *shallow benthics*. In this model, it is unclear what evolutionary forces drove these back introgressions of the non-inverted alleles, but would imply these inversions have not continuously provided the genetic sex determination mechanism since their inception.

The distribution of the inversions in the benthic sub-radiation is uniquely complicated in Lake Malawi. Inverted and non-inverted haplotypes for five of the six inversions are segregating within this group. Additionally, the presence of the 9, 11, and 20 inversions within *benthics* violates the whole genome phylogeny. We propose these three inversions originated within the *Diplotaxodon* group and spread to a very early *benthic* ancestor via introgression. We favor this over incomplete lineage sorting, which would also lead to violations of the phylogeny, for a few reasons. First, the dynamic history of the lake levels is consistent with times of extinction and hybridization^48–50^. Second, genomic analysis of extant cichlids provided evidence for a large amount of hybridization events^36,55^. Finally, the close genetic distance between *Diplotaxodons* and *benthics* that carry the inverted haplotype (**Figure 5**) is consistent with hybridization. Hybridization has been proposed to fuel adaptive radiations of cichlids in all three great African lakes (Malawi, Tanganyika, and Victoria) ^27,28,56^ and our work suggests that an ancient hybridization event between early *benthic* and *Diplotaxodon* ancestors occurred prior to the benthic radiation. It will be interesting to study the role of these three inversions in *benthic* species. *Diplotaxodon* are pelagic, deep-water dwellers that feed on plankton or small fish while *benthics* live near the shore over sandy and muddy habitats^30^. It is not obvious how the traits of the *Diplotaxodon* would benefit *benthic* animals that live in such dissimilar environments. However, the initial habitats that the *benthic* ancestors utilized is not known and potentially their initial habitat was in deeper waters than the *AC* ancestors.

This report provides an initial framework for understanding the role of inversions in the adaptive radiation of Lake Malawi cichlids. Future work identifying and characterizing causal alleles captured within these inversions will enable us to discern the specific mechanisms by which they contribute to speciation using this remarkable evolutionary system as a model for the evolution of all animals.

## ACKNOWLEDGEMENTS

We thank Krish Roy for the acquisition of the Bionano Saphyr system, Manasi Pimpley for assistance in adapting existing Bionano kits to cichlid tissue, and the Molecular Evolution Core Laboratory at the Parker H. Petit Institute for Bioengineering and Bioscience at the Georgia Institute of Technology for the use of their shared equipment, services, and expertise. This work was supported in part by NIH R35 GM139594 and the Nelson and Bennie Abell Professorship to P.T.M., R01GM144560 to J.T.S., and U.S. National Science Foundation (DEB-1830753) to T.D.K..

## METHODS

### Samples

Wild-caught cichlids were acquired from Old World Exotic Fish Inc in 2022 and Cichlidenstadel in 2024. Lab-reared species were purchased from Southeast Cichlids, Old World Exotic Fish Inc., and Cichlidenstadel at various points in the past two decades (**Table S2**). Fish were delivered live to the Georgia Institute of Technology cichlid aquaculture facilities. All samples were collected following anesthetization with tricaine according to procedures approved by the Institutional Animal Care and Use Committee (IACUC protocol numbers A100029 and A100569).

Caudal fins were collected from subjects and flash frozen on powdered dry ice before storage at −80°C. Care was taken to minimize freeze-thaw cycles prior to DNA extraction via Bionano or Qiagen protocols. If necessary, fins were sectioned on a sterile aluminum block set in dry ice prior to digestion.

Blood was collected following rapid decapitation of anesthetized subjects. Wide-bore, low-retention pipette tips, 0.5M EDTA (pH 8.0, Invitrogen Cat# 15575-038), and Bionano Cell Buffer (Part Number 20374) prevented blood from clotting at room temperature. Samples were used immediately for Bionano DNA extraction (see below).

### Bionano

Bionano DNA extractions were predominantly performed on fresh whole blood samples (see above). Blood concentration was determined via a manual logical count performed on a hemocytometer and 2 million cells were carried forward for DNA extraction with the Bionano SP-G2 Blood & Cell Culture DNA Isolation Kit (Part Number 80060) as per the Bionano Prep SP-G2 Fresh Human Blood DNA Isolation Protocol (Document Number CG-00005, Rev C). Because nucleated cichlid blood had high concentrations, steps 3 and 4 were skipped and samples were resuspended in Bionano Cell Buffer to a total volume of 200µL. Following extraction, samples were allowed to homogenize undisturbed at room temperature for at minimum 72 hours prior to quantification using the Qubit dsDNA Quantitation, Broad Range kit (ThermoFisher, Cat # Q32853).

Bionano data for a single *Protomelas taeniolatus* sample (PT_2003_m, see **Table_S2**) was generated from fin tissue using a combination of the Bionano Prep SP Tissue and Tumor DNA Isolation Protocol (Document Number 30339, Rev A) and the Bionano Prep SP-G2 Fresh Human Blood DNA Isolation Protocol.

750ng of homogenized DNA was fluorescently labeled using the Bionano Direct Label and Stain-G2 (DLS- G2) Kit (Part Number 80046, Protocol - CG-30553-1, Rev E). Labeled DNA was quantified using the Qubit dsDNA Quantitation, High Sensitivity kit (ThermoFisher, Cat # Q32854) and loaded onto a flow cell within a Bionano Saphyr Chip G3.3 (Part Number 20440). Samples were imaged on the Bionano Saphyr with at least 500Gbp of data collected per sample.

Molecules for each sample were assembled using the Bionano *de novo* assembly pipeline on Bionano Access Version 1.7. The preassembly setting was turned on, and the variant annotation parameters were deselected for every assembly. *De novo* maps were aligned to the Mzebra_GT3a genome.

### PacBio HiFi sequencing and genome assembly

PacBio *de novo* assemblies were generated from HiFi reads across three sequencing runs. The MZ_GT3.3 and YH_GT1.3 were sequenced following DNA extraction from frozen caudal fin tissue using the Qiagen MagAttract HMW DNA Kit (Cat. No. 67563). In the first sequencing run a *Metriaclima zebra* female was sequenced sequenced on a PacBio Sequel II system by the Georgia Genomics and Bioinformatics core. In the second run, the same the *Metriaclima zebra* female and an *Aulonocara sp. ‘chitande type north’ Nkhata Bay* female was sequenced on a PacBio Revio instrument by the HudsonAlpha Institute for Biotechnology. Reads from both runs were used to assemble the MZ_GT3.3 genome and reads from the second run alone were used to assemble the YH_GT1.3 genome (see details below).

Mzebra_GT3a, MZ_GT3.2, YH_GT1.1, and YH_GT1.2 were assembled using HiFi reads generated with DNA extracted from fresh whole blood and heart tissue. DNA was extracted via the PacBio Nanobind Tissue Kit RT (Part Number 102-208-000) following a modified DNA from animal tissue using the Nanobind® PanDNA kit protocol (Part Number 102-574-600). Blood and heart tissue were combined and homogenized together using a Qiagen TissueRuptor (Qiagen 9002755) and centrifugation g-force was halved during the DNA extraction. DNA fragments <25kb were removed using the PacBio Short Read Elimination kit (Part Number 102-208-300). Library preparation and sequencing was performed by the University of Maryland Institute for Genome Sciences on a PacBio Revio instrument.

HiFi reads from all samples were assembled using the Mabs de novo assembler (v2.28)^57^ using the `mabs-hifiasm` algorithm and default parameters. Each assembly from mabs-hifiasm was evaluated using Inspector (`inspector.py` v1.0.1)^58^ run on default parameters. The assemblies were error corrected with `inspector-correct.py` (v1.0) to resolve large structural errors. These error-corrected contigs from were uploaded to Bionano Access.

The MZ_GT3.2, MZ_GT3.3, YH_GT1.1, YH_GT1.2 and YH_GT1.3 scaffold-level assemblies were generated by bridging gaps in their contigs using Bionano maps via the single-enzyme Bionano Hybrid Scaffold pipeline using default parameters. A unique set of Bionano maps was used to scaffold the contigs from each *de novo* assembly. The resulting hybrid scaffold `NCBI.fasta` file (which denotes gaps filled by Bionano maps with stretches of N nucleotides) was concatenated with any unscaffolded contigs from the error-corrected mabs assembly. These five scaffold-level assemblies have been deposited at DDBJ/ENA/GenBank (accessions TBD).

A scaffold-level genome was identically generated for Mzebra_GT3a. These scaffolds were further aligned to the corrected UMD2a genome (see below) and anchored to linkage groups using D-GENIES^59^ with the Minimap2 v2.28 aligner and “Many repeats” options. An anchored genome was output using the Query assembled as reference export option.

Note that the mitochondrial DNA sequence in Mzebra_GT3a is the same as the mitochondria assembled in UMD2a. This Whole Genome Shotgun project has been deposited at DDBJ/ENA/GenBank under the accession JBEVYI000000000. The version described in this paper is version JBEVYI010000000.

### Correcting errors in M_zebra_UMD2a

We generated Bionano molecules for 3 male and 1 female *Metriaclima zebra*, assembled Bionano maps, and aligned them to the UMD2a reference genome. We reordered or reoriented contigs that were revealed as misassemblies or inversions, respectively, in all four Bionano maps and corresponded to breakpoints between contigs in UMD2a. By filtering interchromosomal translocation calls in the Bionano Access software, we were able to insert unmapped contigs into the established linkage groups.

### Illumina extractions and sequencing

DNA was extracted from fresh or frozen fin tissues for short-read sequencing using the Qiagen MagAttract HMW DNA Kit (Cat. No. 67563) or the Qiagen DNeasy Blood & Tissue Kit (Cat No. 69504) using the Manual Purification of High-Molecular Weight Genomic DNA from Fresh or Frozen Tissue and Purification of Total DNA from Animal Tissues (Spin-Column) protocols respectively. DNA was delivered to the Georgia Tech Molecular Evolution Core where libraries were prepared with the NEBNext® Ultra™ II FS DNA Library Prep Kit for Illumina (NEB #E6177) using the Protocol for FS DNA Library Prep Kit (E7805, E6177) with Inputs ≥100 ng. A small subset of samples was sent to an external collaborator by the Molecular Evolution Core where library preparation was performed using the KAPA HyperPrep Kit (Roche, Material Number: 07962363001). Samples across all runs were sequenced on a NovaSeq 6000 instrument using v1.5 chemistry.

### Variant calling and PCA analysis

We provide a link to a github repository (https://github.com/ptmcgrat/CichlidSRSequencing/tree/Kumar_eLife) containing the scripts used for the major analysis in the paper. Because our data is behind a secure Dropbox account, readers will be limited to seeing the exact programs, filters, and parameters used for this manuscript embedded within each script.

Fastq files from Illumina sequencing were converted to UBAM format using the `gatk`^60^ `FastqToSam` algorithm. UBAM files were used for alignment to Mzebra_GT3 using `bwa`^61^ `mem`; the `-M` and `-p` flags were used. To carry forward metadata from the unaligned BAMs, `gatk MergeBamAlignment` was called on the alignments. BAM files were converted to GVCF format using `gatk HaplotypeCaller`. Variant calling was performed using the GVCF files for the analysis cohort. The `gatk GenomicsDBImport` and `gatk GenotypeGVCFs` algorithms were used to generate a master vcf file which was subsequently filtered with `gatk VariantFiltration` (parameters below). The variants that passed filtering were used for PCA analysis.

PCA was performed with `plink2`^62^. To avoid overrepresenting species from the cohort, a core subset of the samples was used for PC 1 and 2 calculations (**Table S2**, column S). Eigenvectors from the whole cohort were then plotted on this PC space and visualized using `plotly`.

Note that GATK ≥ v4.3.0.0 and python ≥ 3.7 were used for these analyses. Custom python scripts were written to automate and parallelize processing of the samples in the cohorts.

The commands used for variant calling and PCA are the following:

#### UBAM Generation

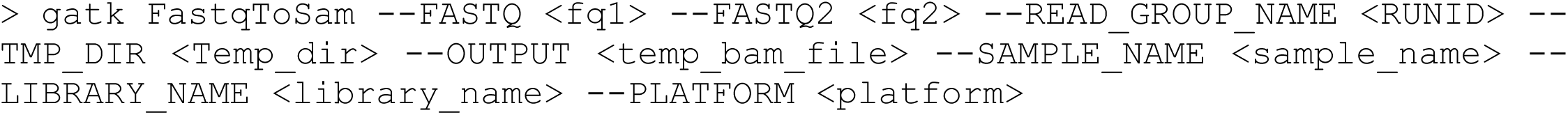

#### Alignment to GT3

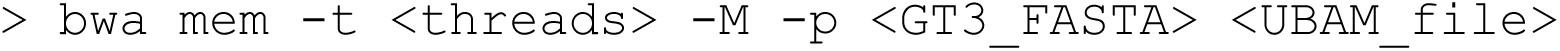

#### Merge metadata from UBAM to BAM

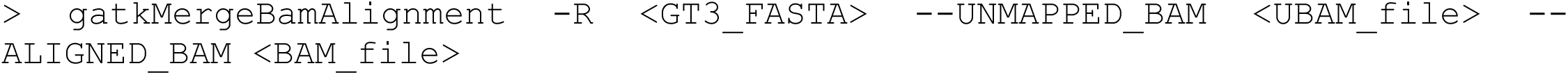

#### Generate GVCFS

`> gatk HaplotypeCaller -R <GT3_FASTA> -I <BAM_FILE> -ERC GVCF -O <GVCF_FILE> -ERC GVCF` is used to generate GVCF files that can be used for subsequent joint genotyping

#### Read Depth Information

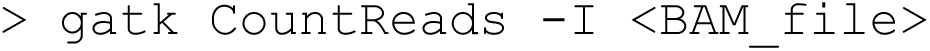

#### Generate GenomicsDB

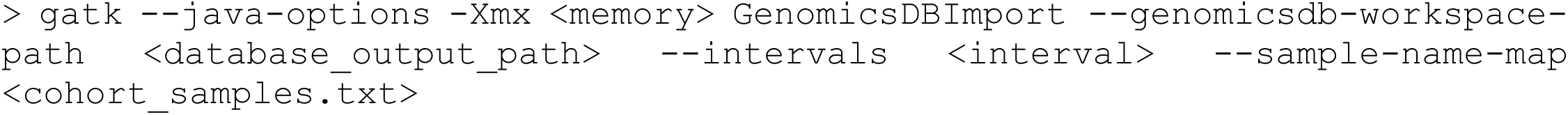

#### Joint Genotyping

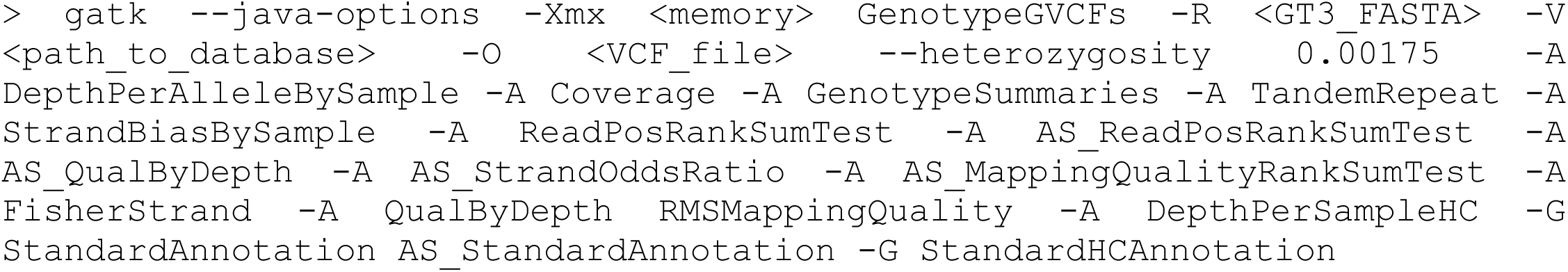

Note that an average measure of heterozygosity, 0.00175 was used from what was reported in Malinksy *et al*.^30^.

#### Filtering Variants

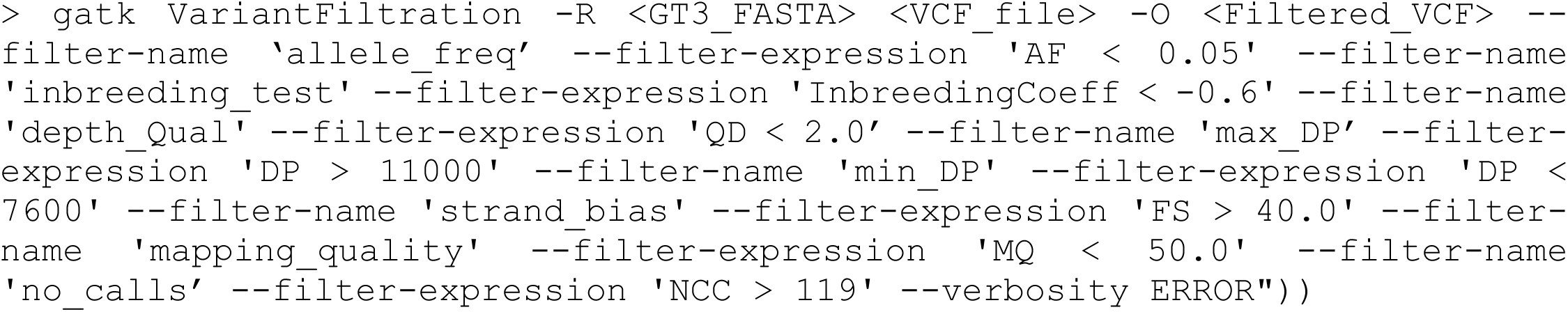

Variants were filtered by allele frequency, inbreeding coefficient, quality by depth, depth, fisher’s exact test for strand bias, mapping quality, and by excess missingness according to methods published in Malinksky *et al*.^30^. We filtered by depth to include variants with a mean depth per sample between 10% and 95% of the distribution of all variant depths.

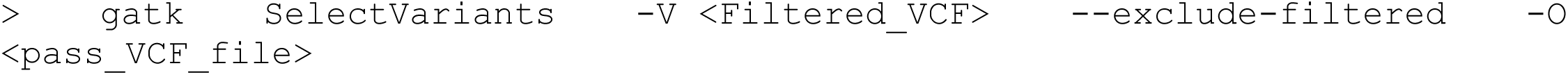

#### PCA

For PCA, first a subset of the pass_VCF_file containing the samples we used to create the principal components was generated using `bcftools`^63^. Next, the relevant pfiles were generated for both the pass_VCF_file and the subset_samples_VCF. Linkage pruning was only performed for the whole genome and whole chromosome PCAs using the –indep-pairwise 50 5 0.1 flag and parameters. Since inversions inherently link together variants, linkage pruning was not used when restricting the analysis to within inverted regions. A linear scoring system is generated using the subset_samples_VCF pfiles. This scoring system is applied to the genotype matrix for the whole cohort’s samples to scale each sample consistently. The resulting first two eigenvectors per sample from the .sscore file are plotted with `plotly`.

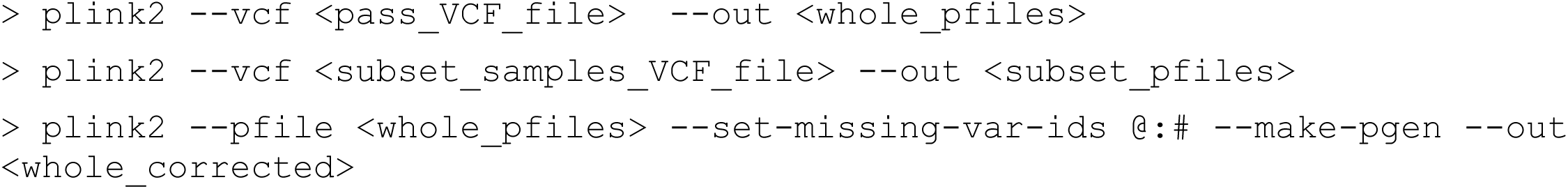

LD Pruning for Whole genome and whole chromosome PCAs:

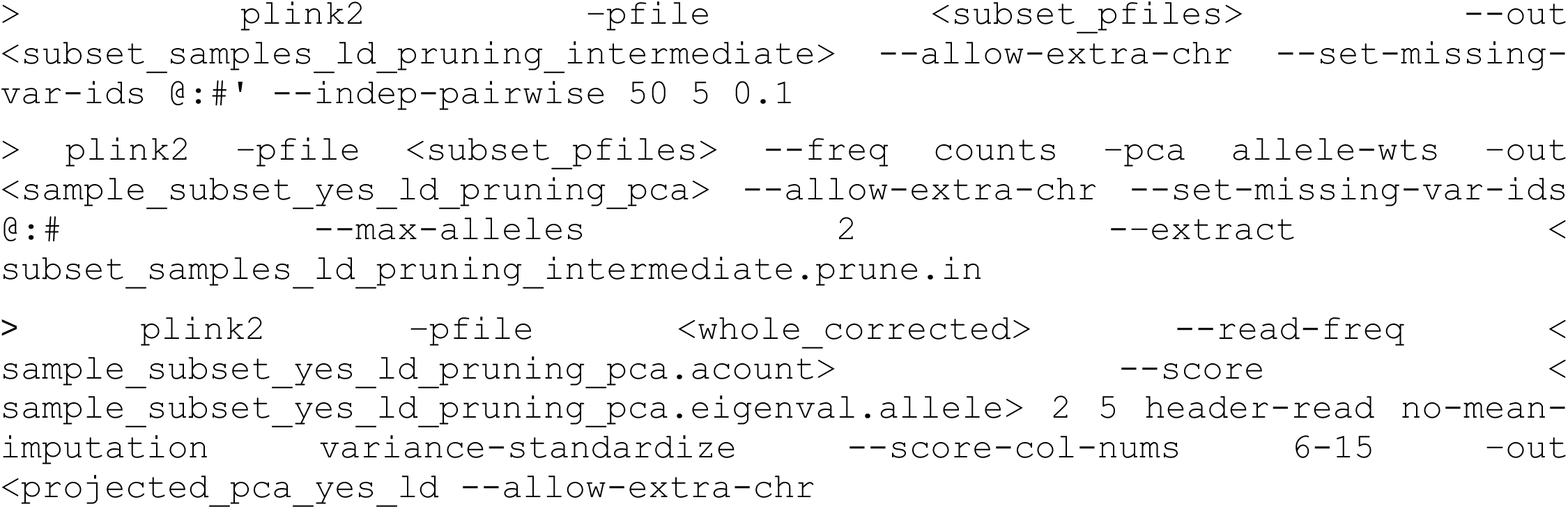

No LD Pruning for inverted region PCAs:

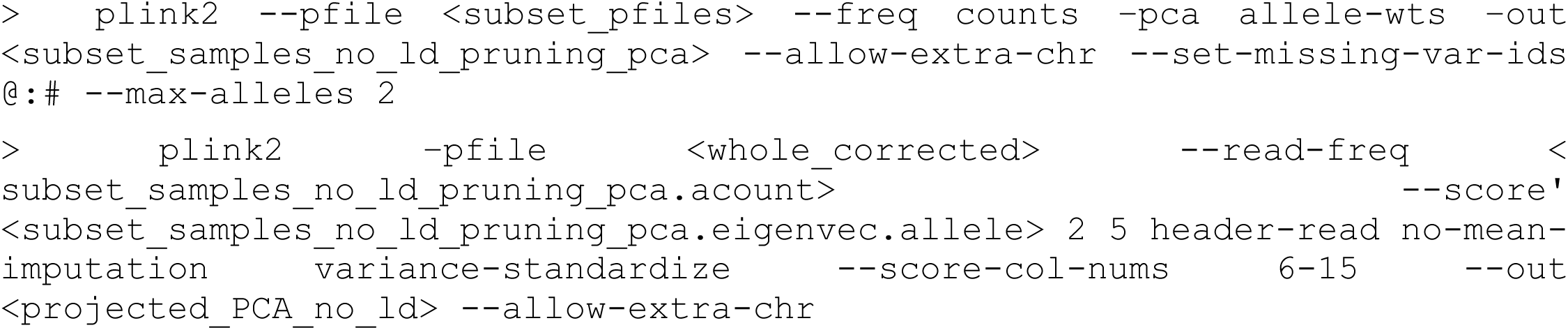

### Genome alignment

Genomes were aligned using `minimap2`^64^ (v2.22) to align the query genome to the M_zebra_GT3 reference using default settings. Custom scripts were used to convert the output paf files to dataframe objects, filtering removed alignments that were 1) secondary matches, 2) had less than 30% percent identify, and 3) were less than 10,000 base pairs in length. Further, we excluded any alignments from a contig containing less than 2,000,000 total bp of alignments, to account for repetitive DNA. Alignments were plotted using `seaborn`. We estimated the intervals containing the breakpoints for the inversions using genome alignments of Aulonocara sp. ‘chitande type north’ Nkhata Bay and M_zebra_GT3 (**Table S4**).

### Phylogenies

To create phylogenies, `bcftools` was used to filter out non SNV variants from the master vcf file. `Bcftools` was also used to create individual vcf files for each inversion, filtering out SNVs that fell outside of the inverted region. `Vcf2philip`^65^ was used to convert the to the phylip format, using the -m 50 option to filter out SNVs with less than 50 genotyped samples. `Iqtree2`^66^ was used to create trees and estimate confidence values for each node using the following options: `-nt’,‘24’,‘-mem’,‘96G’,‘- v’,‘--seqtype’, ‘DNA’,‘-m’,‘GTR+I+G’,‘-B’,‘1000’`. Trees were visualized with `iTol`^67^ or `ete3`^68^.

### Nucleotide diversity analysis

To analyze the Pixy between individuals of different species, we used `scikit-allel` (v1.3.8)^69^ to calculate the genetic difference between each sample using the `allel.pairwise_distance` function and the `citblock` for the metric. We restricted the analysis to benthic animals that carried the inversion and compared these to animals of various ecogroups. We used the `seaborn` function `hist plot` to create density histograms for each of the inversions.

### Fst & pedigree

A vcf file was created for the *Aulonocara* samples listed in **Table S2** (Column I). The PCA approach (described above) was used to genotype each of the samples for the inversion on 10. For the F_ST_ analysis, we used `scikit-allel` (v1.3.8) to perform `allel.average_weir_cockerham_fst` with a window size of 100. For the heterozygosity analysis, we used the `allel.heterozygosity_observed` function on both the male and female population, subtracting the average heterozygosity from the male offspring from the female offspring.

## DATA AVAILABILITY

All Illumina and PacBio sequencing reads have been deposited to the NCBI Short Read archive at BioProject PRJNA1112855. The VCF files used in these analyses are available through the Dryad Digital Repository (TBD).

