## Supplementary material for "Large inversions in Lake Malawi cichlids are associated with habitat preference, lineage, and sex determination": FigureS1_HeterozygousIndividuals.pdf

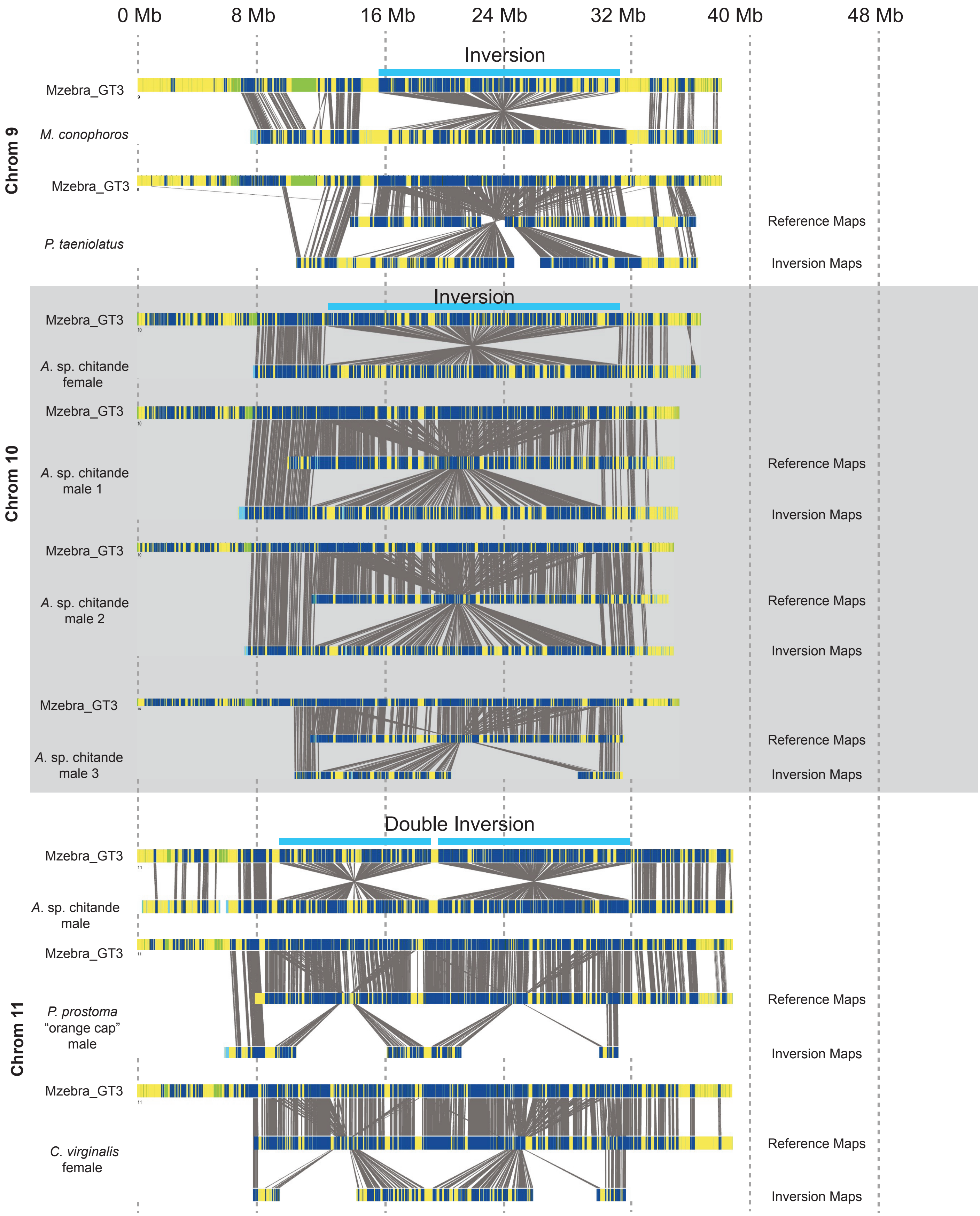


**Figure S1 - Bionano** maps from individuals that were heterozygous for the inversions on chromosomes 9, 10, or 13. Bionano maps that support the non-inverted haplotype are shown in the middle while Bionano maps that support the inverted haplotype are shown below.
