## Supplementary material for "Large inversions in Lake Malawi cichlids are associated with habitat preference, lineage, and sex determination": FigureS5_LG11_Alignment.pdf

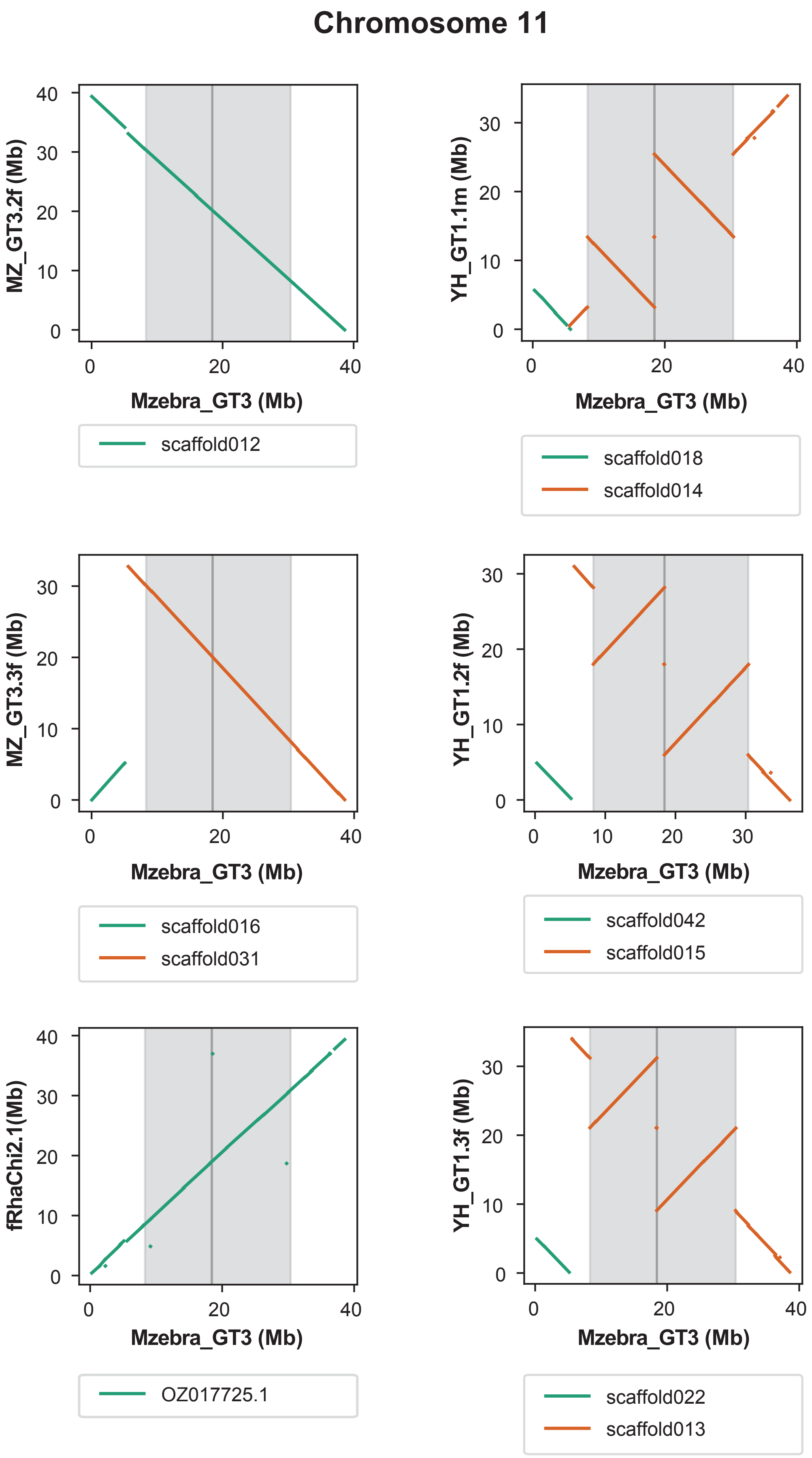


**Figure S5 - Genome alignments for chromosome 11**. Dot plots comparing the reference M_zebra_GT3a (x-axis) to six other genomes created from Lake Malawi cichlid fish (y-axis). Alignments are colored by scaffold names (see legends). MZGT3.2f and MZGT3.3f were each created from unique Metriaclima zebra female fish. YH_GT1.1f, YH_GT1.2m, and YH_GT1.3f were created from a single female, male, and female Aulonocara sp. ‘chitande type north’ Nkhata Bay fish, respectively. fRhaChi2.1 was a previously published genome generated from a Rhamphochromis sp. chilingali. Grey bar indicates location of the inversion.
