## Supplementary material for "Large inversions in Lake Malawi cichlids are associated with habitat preference, lineage, and sex determination": FigureS8_InversionAncestry.pdf

### Outgroup alignments

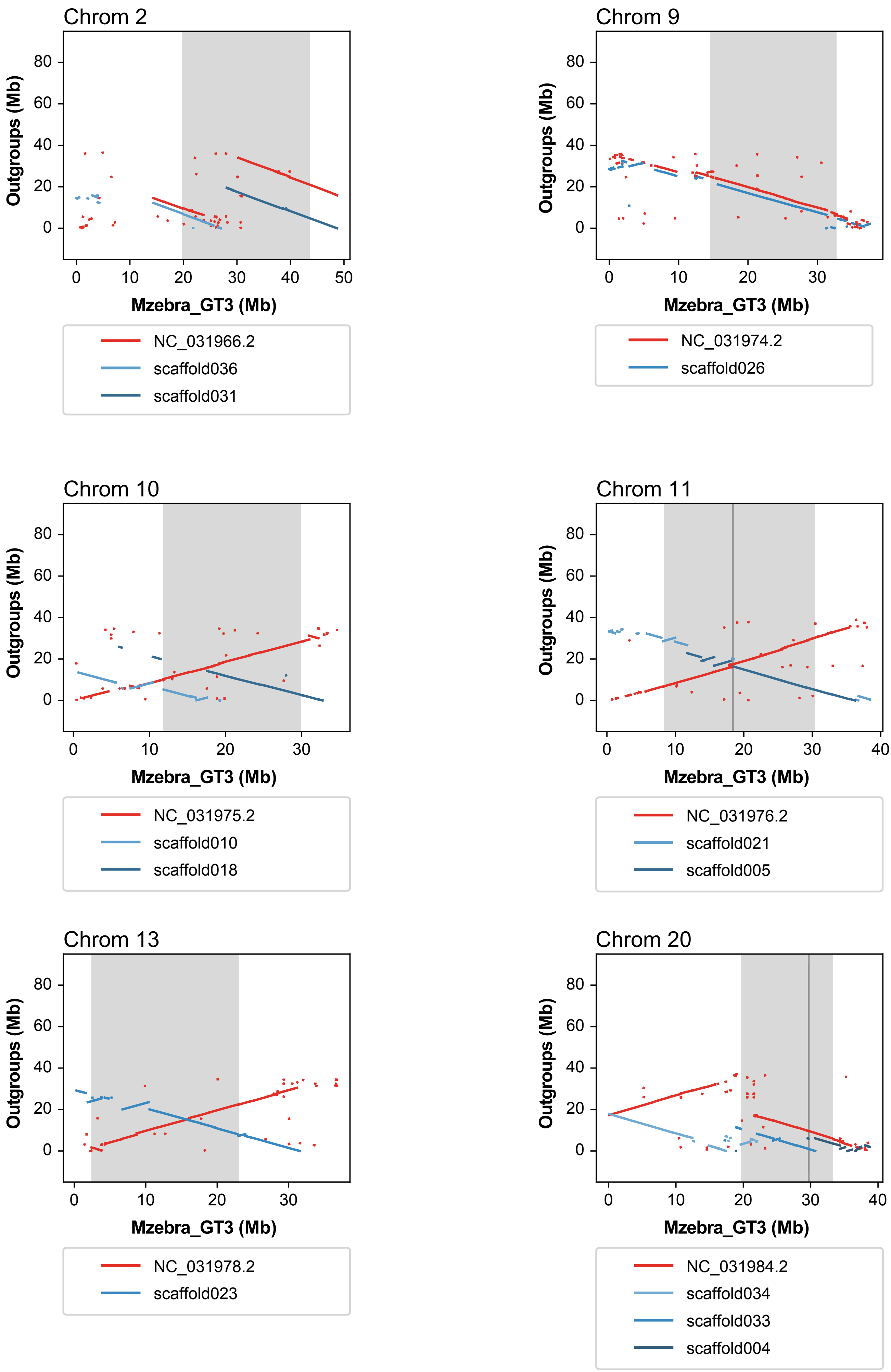


**Figure S8 - Genome alignments to Lake Malawi outgroups**. Dot plots comparing the reference M_zebra_GT3a (x-axis) to Oreochromis nilticos (red) or Pudimilla nyererei (shades of blue) (y-axis). Inversion location listed above each plot. Grey bar indicates location of the inversions and dark gray bars indicate the location of tandem inversion breakpoints.
