## Supplementary material for "Large inversions in Lake Malawi cichlids are associated with habitat preference, lineage, and sex determination": FigureS19.pdf

Chromosome 2

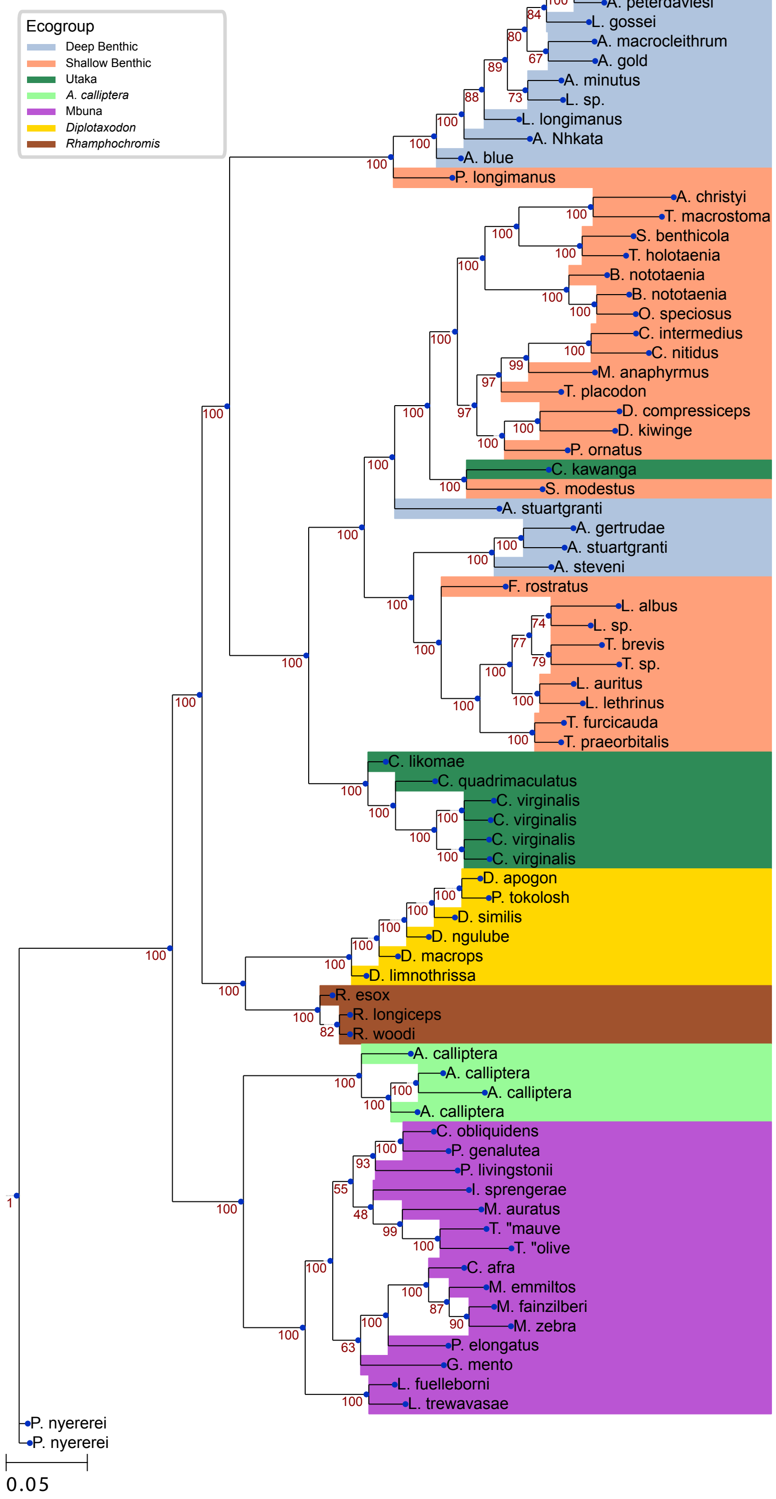


**Figure S19 - Chromosome 2 phylogeny for Lake Malawi cichlids.** A maximum likelihood tree was created from SNV data across the seven major ecogroups of Lake Malawi cichlids, restricted to the inversion region on chromosome 2. Bootstrap support values are shown below each node. Branch lengths are proportional to the number of SNVs per basepair.
