## Supplementary material for "Large inversions in Lake Malawi cichlids are associated with habitat preference, lineage, and sex determination": FigureS21.pdf

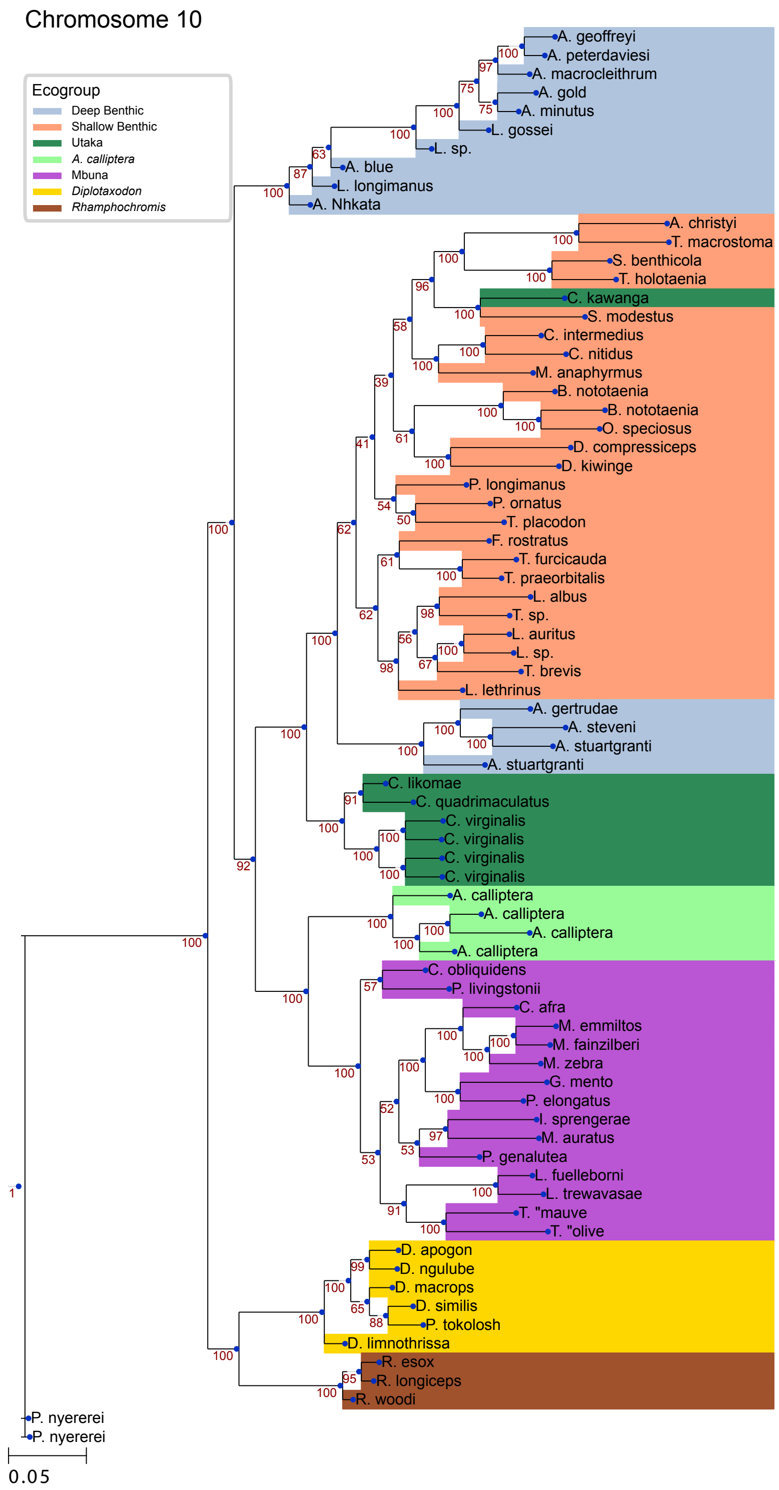


**Figure S21 - Chromosome 10 phylogeny for Lake Malawi cichlids.** A maximum likelihood tree was created from SNV data across the seven major ecogroups of Lake Malawi cichlids, restricted to the inversion region on chromosome 10. Bootstrap support values are shown below each node. Branch lengths are proportional to the number of SNVs per basepair.
