## Supplementary material for "Large inversions in Lake Malawi cichlids are associated with habitat preference, lineage, and sex determination": FigureS23.pdf

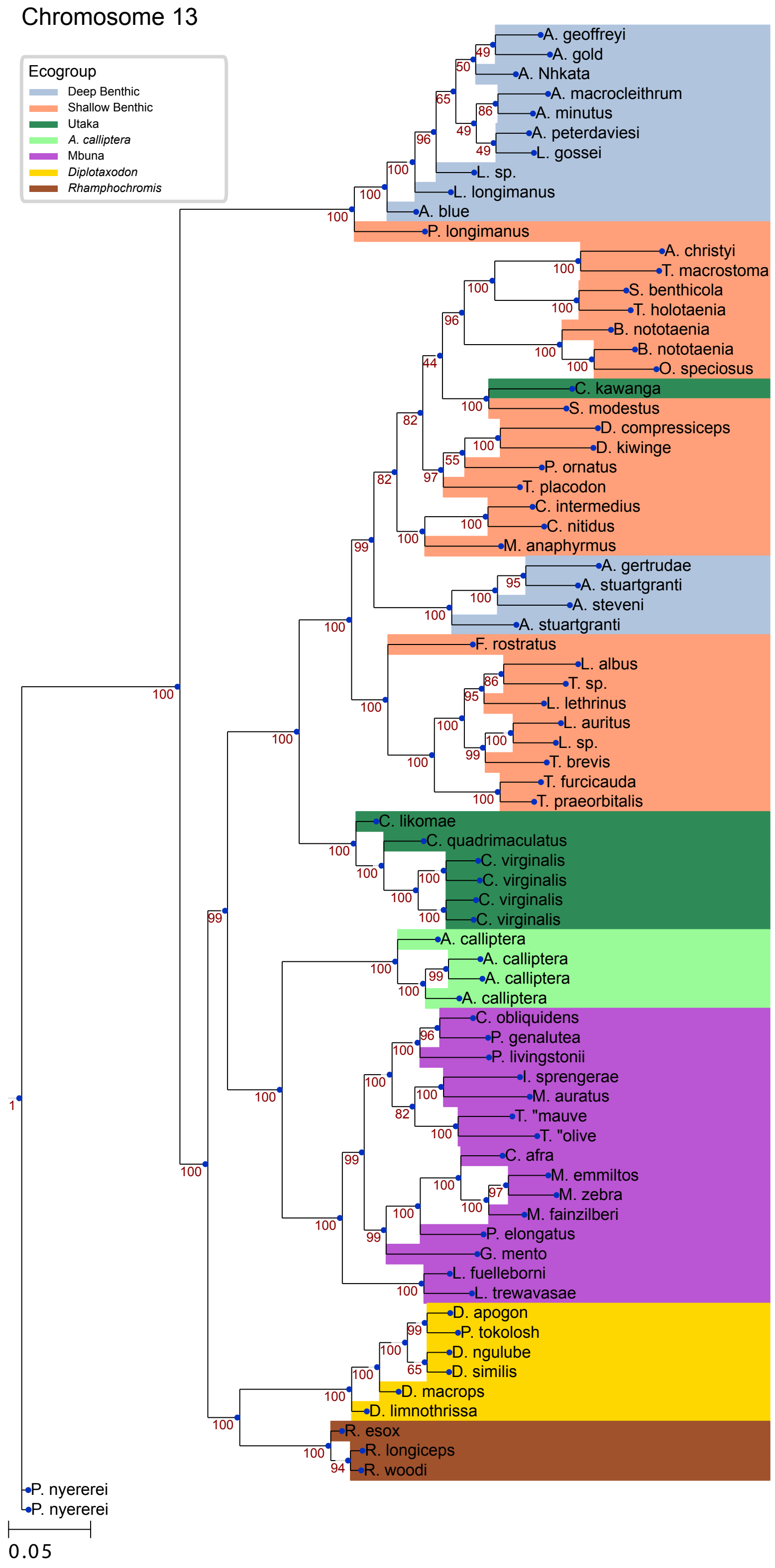


**Figure S23- Chromosome 13 phylogeny for Lake Malawi cichlids.** A maximum likelihood tree was created from SNV data across the seven major ecogroups of Lake Malawi cichlids, restricted to the inversion region on chromosome 13. Bootstrap support values are shown below each node. Branch lengths are proportional to the number of SNVs per basepair.
