## Supplementary material for "Large inversions in Lake Malawi cichlids are associated with habitat preference, lineage, and sex determination": FigureS24.pdf

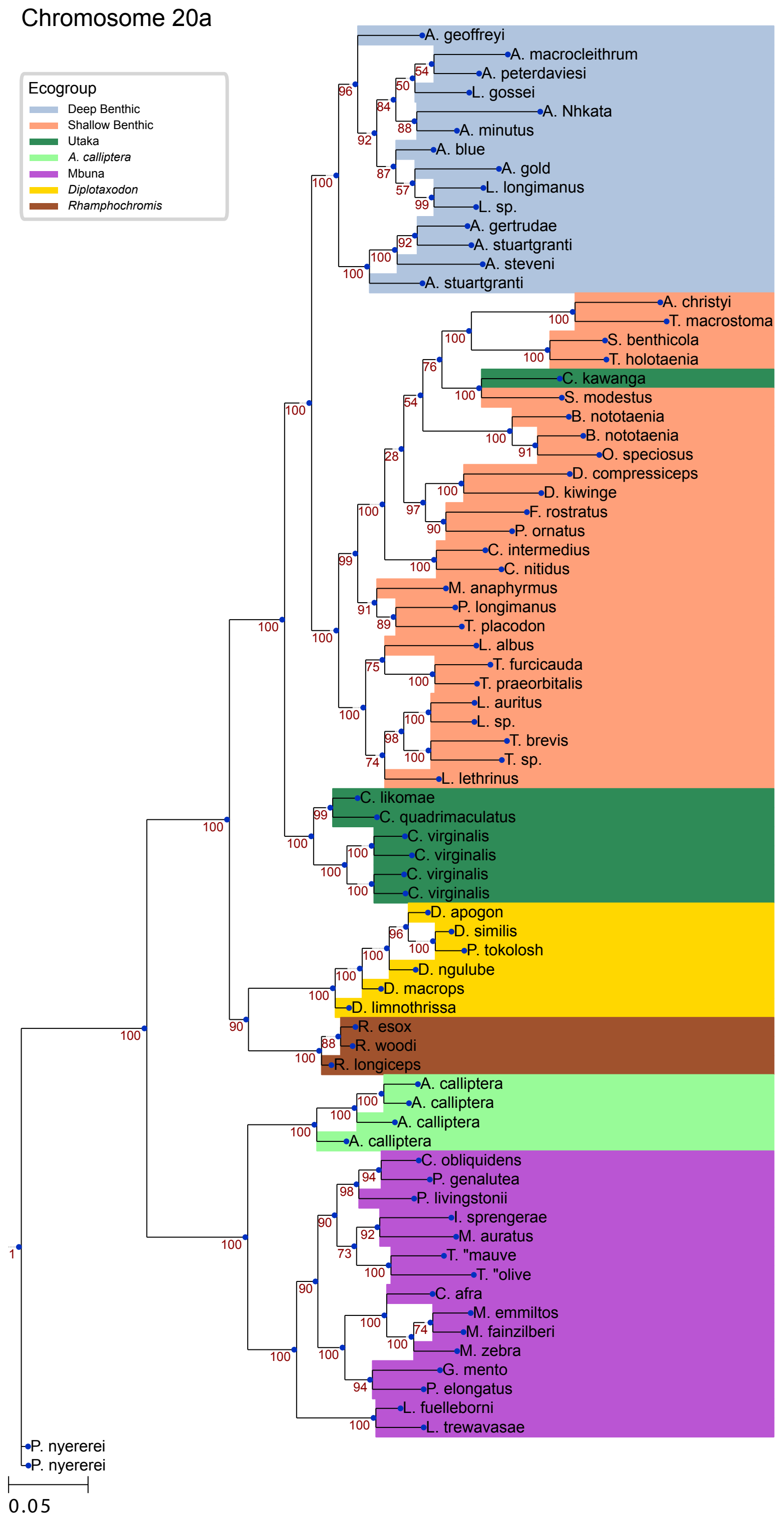


**Figure S24- Chromosome 20a phylogeny for Lake Malawi cichlids.** A maximum likelihood tree was created from SNV data across the seven major ecogroups of Lake Malawi cichlids, restricted to the left-side inversion region on chromosome 20 . Bootstrap support values are shown below each node. Branch lengths are proportional to the number of SNVs per basepair.
