## Supplementary material for "Large inversions in Lake Malawi cichlids are associated with habitat preference, lineage, and sex determination": TableS1_Reference Comparison.docx

| **Table 1** |  | |  |
| --- | --- | --- | --- |
| **LG** | **UMD2a** | **GT3a** | |
| 1 | 38,676,823 | | 45,605,991 |
| 2 | 32,660,920 | | 48,807,247 |
| 3 | 37,314,939 | | 64,350,661 |
| 4 | 30,518,969 | | 37,413,276 |
| 5 | 36,170,306 | | 42,954,235 |
| 6 | 39,774,503 | | 45,268,537 |
| 7 | 64,916,660 | | 69,537,369 |
| 8 | 23,971,387 | | 31,062,834 |
| 9 | 21,023,328 | | 37,905,347 |
| 10 | 32,356,376 | | 34,949,860 |
| 11 | 32,446,080 | | 38,663,802 |
| 12 | 34,086,040 | | 40,266,177 |
| 13 | 32,072,427 | | 36,767,363 |
| 14 | 37,870,038 | | 40,972,746 |
| 15 | 34,548,429 | | 43,150,661 |
| 16 | 34,742,138 | | 37,418,162 |
| 17 | 35,776,103 | | 40,684,106 |
| 18 | 29,505,383 | | 38,717,768 |
| 19 | 25,963,618 | | 32,005,253 |
| 20 | 29,789,237 | | 38,814,023 |
| 22 | 34,725,690 | | 38,466,051 |
| 23 | 42,088,218 | | 50,161,205 |
| **DNA placed on LGs (bp)** | **760,997,612** | | **933,942,674** |
| Unplaced contigs (#) | 1667 | | 41 |
| Unplaced contigs (bp) | 196,471,068 | | 28,232,709 |
| mtDNA | 16,582 | | 16,582 |
| **Genome size (bp)** | **957,485,262** | | **962,191,965** |
