## Supplementary material for "Large inversions in Lake Malawi cichlids are associated with habitat preference, lineage, and sex determination": TableS3_GenomeAssemblies.docx

| **Table S3** |  |  |  |
| --- | --- | --- | --- |
| **Assembly** | **# Scaffolds** | **Size** | **N50** |
| MZ_GT3.2 | 215 | 959,221,546 | 32,733,271 |
| MZ_GT3.3 | 198 | 973,186,388 | 34,500,616 |
| YH_GT1.1 | 199 | 972,426,806 | 34,006,260 |
| YH_GT1.2 | 191 | 976,497,133 | 35,391,406 |
| YH_GT1.3 | 321 | 959,719,201 | 28,082,466 |
