## Supplementary material for "Large inversions in Lake Malawi cichlids are associated with habitat preference, lineage, and sex determination": TableS4_BreakpointEstimates.docx

| **Table S4** |  |  |  |  |
| --- | --- | --- | --- | --- |
| **Chromosome** | **Breakpoint 1 Interval Start** | **Breakpoint 1 Interval End** | **Breakpoint 2 Interval Start** | **Breakpoint 2 Interval End** |
| 2 | 19743639 | 20311160 | 43254805 | 43658853 |
| 9 | 14453796 | 15649299 | 32255605 | 33496468 |
| 10 | 11674905 | 11855817 | 29898615 | 29898615 |
| 11a | 8302039 | 8309764 | 18425216 | 18425216 |
| 11b | 18425216 | 18425216 | 30371888 | 30459686 |
| 13 | 2249541 | 2453698 | 23046928 | 23131968 |
| 20a | 19614379 | 19689710 | 29716509 | 29673642 |
| 20b | 29716509 | 29673642 | 32872827 | 33764042 |
